# Characterization of PROTACs by Ternary Complex Landscape Exploration using Monte Carlo simulations

**DOI:** 10.1101/2025.04.11.648328

**Authors:** Anna M. Diaz-Rovira, Carles Perez-Lopez, Jordi Martín-Pérez, Chiara Pallara, Lucía Díaz, Victor Guallar

**Affiliations:** Barcelona Supercomputing Center, Barcelona, Spain; Doctoral program in Theoretical Chemistry and Computational Modelling, Barcelona University, Spain; Nostrum Biodiscovery S.L., Barcelona, Spain; Institució Catalana de Recerca i Estudis Avançats (ICREA), Barcelona, Spain

**Keywords:** PROTAC, Energy Landscape Exploration, Monte Carlo, Protein-Protein Interactions, Targeted Protein Degradation

## Abstract

PROTACs (Proteolysis-Targeting Chimeras) have emerged as a powerful modality for targeted protein degradation, yet their optimization still relies heavily on trial-and-error methods. A key factor in PROTAC degradation efficiency is the formation of the ternary complex (TC) between the PROTAC and its target proteins. However, due to their dynamic nature, PROTAC-mediated TCs can adopt multiple conformations, making their characterization challenging. Computational methods that account for this flexibility can provide more accurate predictions aligned with experimental results. Here, we explore the dynamic nature of TCs by analyzing their energy landscapes using protein-protein docking coupled with Monte Carlo sampling. This approach enables the identification of energetically relevant TC conformations, including those observed in experimental crystal structures, and allows estimation of thermodynamic and kinetic stability, as shown for a set of VHL-WDR5 PROTACs. Insights from these landscapes could support the screening and optimization of tens of similar PROTACs based on TC stability.

## Introduction

Proximity-inducing pharmacology has transformed therapeutic strategies to tackle undruggable targets. In the field of targeted protein degradation (TPD), PROteolysis TArgeting Chimeras (PROTACs) have emerged as a promising alternative modality, with several candidates advancing to clinical trials as alternative therapeutic modalities.^1–4^ Their mechanism of action relies on inducing proximity between the E3 ubiquitin ligase and the protein of interest (POI), thus promoting POI ubiquitination and subsequent degradation via the ubiquitin-proteasome system.^5,6^ Therefore, PROTAC molecules are designed to simultaneously bind both the E3 ligase and the POI through two ligand moieties (commonly referred to as warheads) connected by a chemical linker. The interaction of the warheads with their respective targets promotes the formation of the ternary complex (TC), which is crucial for the POI ubiquitination and degradation.

Given the central role of the TC formation for efficient degradation, its structural characterization is of great interest. However, capturing these structures is particularly challenging due to their massive size, complexity, and dynamic nature.^7,8^ X-ray crystallography provides high-resolution snapshots of TC conformations, but growing evidence suggests that static structures might be biased to crystallization conditions and often fail to capture the full range of dynamic conformations present in PROTAC-mediated TCs.^7,9–11^ This effect could be even more drastic in weak and/or flexible protein-protein interactions. In fact, multiple configurations can coexist for a given E3 ligase–POI pair, as reported by multiple systems.^7,8,10,12^ The different conformations may exhibit different degrees of stability and functionality, with some configurations potentially representing transient or nonproductive states (not leading to ubiquitination, for example). Consequently, the observable TC stability—measured as ternary dissociation constant (K_D_)—reflects the combined contribution of all possible conformations. This structural heterogeneity highlights the need for computational tools that can exhaustively explore TC energy landscapes and link diverse conformations to their functional outcomes.

Despite significant advances in artificial intelligence-based protein structure prediction methods like AlphaFold2,^13^ these algorithms often struggle to accurately model TCs,^14^ particularly at flexible and dynamic interfaces.^15^ In these regards, methods that rely on co-evolutionary signals may lack sufficient information to predict interactions between proteins that do not naturally associate, a common scenario in PROTAC-mediated systems. By incorporating small-molecules into the protein folding predictions, new algorithms like AlphaFold3,^16^ significantly improve the model quality of PROTAC-mediated TCs.^17^ However, these models still produce only a static conformation of the TC, failing to capture its inherent flexibility. Consequently, computational modeling remains essential for understanding TC formation and dynamics. While multiple reviews have examined computational approaches such as molecular docking, and molecular dynamics (MD) simulations,^18–21^ challenges remain in fully capturing TC dynamics, highlighting the need for further methods exploration and improvement.

Pioneering computational approaches, such as those by Drummond et al.,^22,23^ focused on Protein-Protein Docking (PPD) and linker sampling, but were limited by their reliance on static structures. Subsequent advancements, including RosettaDock-based protocols,^7,24^ and ensemble-based methods like PRosettaC,^25^ improved accuracy but still struggled to capture the full spectrum of TC dynamics. More recent hybrid strategies, such as MD simulations in conjunction with structural experimental data^9^ or with molecular mechanics with generalized Born and surface area solvation (MM/GBSA) calculations,^26^ have begun to address this limitation. While these methods offer improved insight into TC flexibility, they are often computationally expensive and sometimes require experimental structural data to guide simulations. Alternatively, more computationally efficient techniques, such as Monte Carlo-based energy mapping,^27^ have also been explored to capture the TC landscape. However, these studies do not examine the relation of this energy landscape with experimental stability data of the TC.

In this study, we aim to develop a computational workflow that exhaustively explores the energy landscape of these dynamic TCs, with the ultimate goal of ranking PROTACs based on their ability to form a potential ensemble of stable TCs—a critical factor in the PROTAC design process (**Figure 1**). To achieve this, we integrate PPD with PELE (Protein Energy Landscape Exploration),^28^ a semi-flexible Monte Carlo (MC) sampling technique. In this framework, PPD is used to generate an ensemble of potential protein-protein binding modes, as in previously reported methods. Additionally, herein we compare the PPD sampling when including the PROTAC in the docking (holo mode) or without considering it (apo mode). We then leverage high-performance computing to evaluate the resulting TC energy landscape for each PROTAC using PELE. PELE iteratively explores the energy landscape in a computationally efficient manner by applying small translations and rotations of the ligand protein, PROTAC and side-chain sampling, and minimization steps to refine the potential binding modes provided by the PPD. We first validated the workflow ability to retrieve crystal TC in a set of four PROTACs with differing linkers. Subsequently, we evaluated how the sampled TC landscapes correlate with both TC thermodynamic and kinetic stability of TCs.

**Figure 1.**
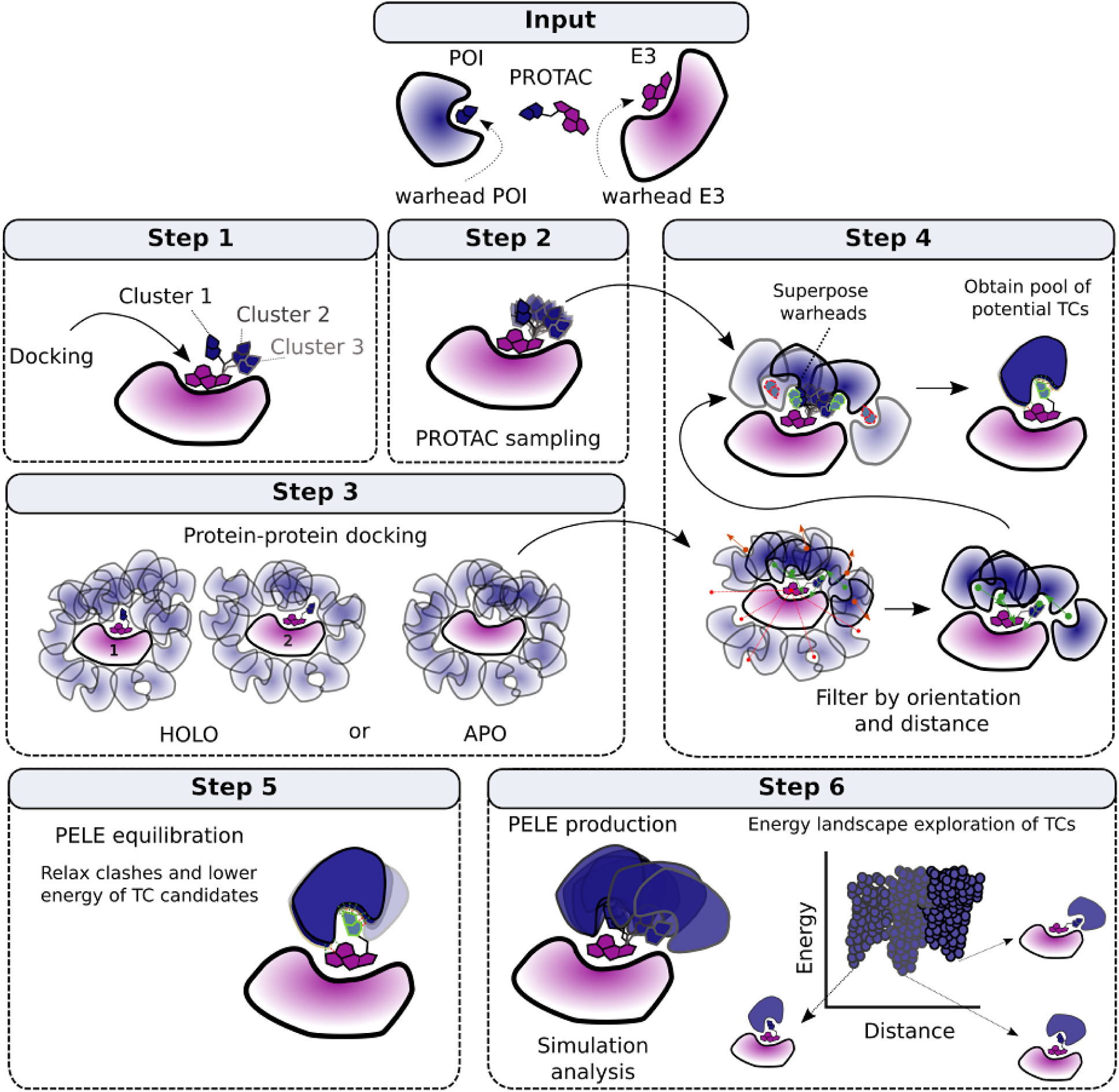
Workflow to study PROTAC TC energy landscape. (Input) The workflow requires as inputs: the POI, the E3 ligase and the PROTAC with known warheads binding site. **(Step 1)** Ligand docking and clustering based on their occupancy volume. **(Step 2)** PROTAC conformational sampling. **(Step 3)** PPD Pose Generation on holo (with PROTAC) **or** apo mode (without PROTAC). **(Step 4)** TC candidate generation derived from filtered PPD poses (Step 3) that meet orientation and spatial constraints for TC formation. Red lines are poses out of the distance, red arrows represent wrong orientations, and the green ones indicate correct structures. Remaining protein poses are paired with PROTAC conformations from Step 2, keeping only those that display correct orientations of the free warhead (green stroke). **(Step 5)** PELE equilibration. **(Step 6)** PELE production and simulation analysis.

## Computational approach

The methodology comprises the following six steps, as outlined in **Figure 1**.

### Workflow input

The workflow takes as input the chemical structure of the PROTACs, the structures of the protein pair (POI and E3 ligase), and information about the warheads’ binding sites and binding modes on their respective recruiting proteins.

### Step 1: Docking the most solvent-exposed PROTAC’s warhead to its recruiting protein

For each protein pair, the warhead is initially docked onto the most solvent-exposed binding site (BS). Since the binding mode of this warhead is typically known, we constraint its pose during the docking and keep it fixed throughout the workflow. This fixed warhead serves as an anchor, allowing the docking conformational sampling to focus on the linker and the more buried (or free) warhead.

Fixing the solvent-exposed BS also offers practical advantages for TC modeling. These sites often involve more flexible or ambiguous protein–ligand contacts, which are difficult to model accurately. Thus, by fixing the exposed warhead, we reduce the system’s degrees of freedom, allowing the more buried (free) warhead to accommodate to its binding site efficiently and naturally—guided by inherent properties such as shape complementarity—without requiring manually defined restraints.

From docking, we generate around 20 poses per PROTAC, which are clustered based on volume to yield a representative set of conformations (typically 1–10). This selection captures a range of linker and free warhead orientations, which is crucial for exploring TC formation in later steps.

### Step 2: PROTAC’s conformational sampling within the binary complex

Using the set of binary complexes obtained in the previous step, we employ our in-house software, PELE, to sample PROTAC conformations within the context of the binary complex, specifically with the least buried warhead already bound. This approach ensures that PROTAC conformations are sampled in a protein-relevant context, effectively avoiding the exploration of conformations that are unlikely to contribute to the formation of the TC from this state.

### Step 3: PPD using FTDock

In parallel with Step 2, we use FTDock^29^ to generate a diverse ensemble of PPD conformations. Within our workflow framework, this process can be performed in two distinct modes:

**i) Holo mode:** This PROTAC-specific approach involves running a PPD for each representative binary complex from Step 1, with the PROTAC embedded in the receptor protein. Retrieving multiple binary complex conformations from the Step 1 is crucial to not impair key protein-protein contacts during the PPD. Each docking generates 92,400 poses, simulating interactions between the binary complex and the free protein to closely mimic the real molecular phenomenon of the TC formation.
**ii) Apo mode:** This PROTAC-independent approach requires only one PPD per protein pair. It assumes that the two proteins have the ability to interact, at least minimally, in their native state, and relies on exhaustive sampling by the PPD software to capture these transient interactions. Compared to the holo mode, this method is more computationally efficient, as it requires just a single PPD run per protein pair.

### Step 4: Generation of TCs candidates

Before generating TC candidates, an initial filtering step is applied to reduce the PPD search space by eliminating docking poses that are unlikely to support viable TC formation. Specifically, PPD poses where the protein binding sites are not aligned face-to-face or are separated by excessive distances (i.e., over 30 Å) are discarded (**Figure S1**). This ensures that only the most relevant PPD poses are considered in the subsequent steps.

To come up with a pool of diverse TC candidates to explore the landscape, we combine the information from the PROTAC sampling conformations from Step 2 and the filtered PPD poses. In particular, we superpose the fixed warhead of each PROTAC conformation to each PPD pose and compute the free warhead RMSD to its known binding mode using three representative atoms for each pose (see Methods, **Figure S2**). We consider that a PPD pose and PROTAC conformation is resulting in a potential TC candidate if the free warhead RMSD is below 2 Å. When this compatibility criterion is met, we refer to the corresponding PPD as a fished PPD and the resulting TC as a fished TC. Finally, for each unique fished PPD we select the TC candidate with the lowest free warhead RMSD to advance to the equilibration step.

### Step 5: Equilibration of TCs candidates with PELE

Most of the previously generated TC candidates have steric clashes between the PROTAC and the free protein. To relieve these clashes and identify low-energy candidates, we use PELE for local refinement. This involves a brief simulation where we apply soft random perturbations to the free protein, followed by small random rotations to the linker and free warhead rotamers (see Methods). Since many candidates may only reach a low-energy state by completely displacing the free warhead from its BS, we filter out those whose free warhead is more than 12 Å away from the BS (see Methods). From the remaining pool of candidates, we select the lowest total energy TC candidate for each unique PPD pose to maintain conformational diversity. Finally, the best 50% of TC candidates in total energy are advanced to production, ensuring that the next step exhaustively samples the most energetically relevant conformations.

### Step 6: TC landscape exploration with PELE

To comprehensively explore the TC landscape, we conduct extended PELE simulations starting from the selected equilibrated TCs (Step 5). We employ AdaptivePELE,^30^ which allows directing the sampling towards TC conformations after doing several unconstrained PELE steps. Specifically, each AdaptivePELE epoch consists of eight PELE steps. Similarly as in the equilibration phase, in each MC step we apply random perturbations to the free protein and randomly sample the PROTAC linker and warhead rotamers. At the end of each epoch, the conformations explored during that epoch are clustered using the PROTAC contact map. Representative structures with the lowest distances between the free warhead in its BS (typically described by hydrogen bond interactions, when possible) are prioritized as starting structures for the next epoch. This approach ensures that, if the free warhead moves away from the BS during sampling, subsequent epochs start from points closer to the BS, improving sampling on the formation of TCs.

Finally, we extract the TC energy landscapes by filtering out all the conformations that are not forming a TC mediated by the PROTAC, that is when the free warhead is not embedded in its BS. Energy distributions from the remaining TC are then analyzed for potential correlation with the experimental ternary K_D_, in order to address the hypothesis that to correlate with experimental ternary K_D_, we need to consider the dynamics of the TC.

## Results and discussion

While many computational approaches aim to recover crystal-like TC conformations,^9,22,25,31^ only a few methods attempt to correlate computational modeling of TCs with TC stability or degradation efficiency.^23,24,32^ These correlations are often solely based on filtering and counting the number of TCs that the PROTAC can form, without considering the energy profile of those complexes. Additionally, most methodologies are benchmarked on PROTAC datasets where degradation efficiency increases with longer linkers,^7,33,34^ but the opposite trend is rarely tested, potentially masking length and flexibility related biases.

In this study, we sought to find correlations between TC landscapes and the experimental TC kinetics and thermodynamics by exhaustively sampling the conformational ensemble of the TCs. Notice that since we solely focus on generating TC landscapes, we did not attempt to correlate with degradation efficiency, which involves additional biochemical factors beyond TC formation.^35^

We benchmarked our workflow on a comprehensive dataset of four PROTACs targeting WDR5 with the VHL E3 ligase,^10,36^ for which crystal structures, ternary K_D_ values, estimated TC association times, TC half-life, and degradation efficiencies are available, enabling us to study both the structural features of the system and its observed TC stability from both the thermodynamics and kinetics perspectives. Interestingly, these PROTACs present varying linker lengths, with the compound with the most thermodynamic stable TC (ms67) having the shortest linker, while the PROTAC forming the least stable TC (8f) is only one carbon shorter than the moderately stable TC mediated by the compound 8g (**Figure 2**). Therefore, this dataset allows us to test our approach on a system that exhibits a complex pattern, with a trend of increased TC stability with longer linkers and an additional one where shorter linkers yield higher TC stability.

**Figure 2.**
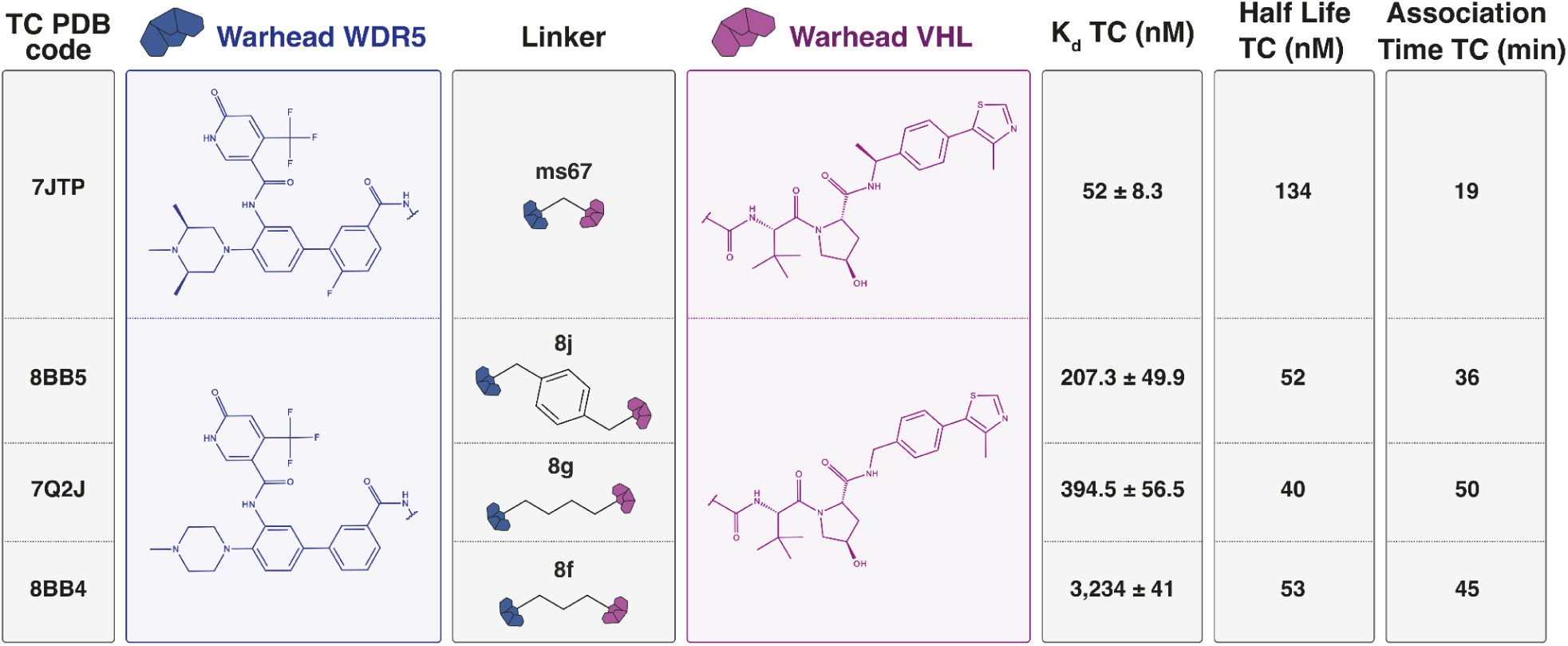
Dataset of PROTACs targeting WDR5 via VHL. These compounds were designed and characterized by Schwalm et al.,^10^ Dölle et al.^36^ and Yu et al..^37^ The experimental metrics were obtained *in cellulo* using the NanoBRET technique. The ternary K_D_ was measured by adding the PROTAC in the presence of both target proteins The estimated TC association time represents the time required to detect the TC signal after PROTAC addition. The TC half-life refers to the time during which the PROTAC can maintain the TC signal upon its formation.

### Structural validation of the workflow

To gain insight into the structural determinants underlying the TC landscape, we first examined the conformational ensembles generated by our workflow. A key validation step is determining whether crystal-like conformations can be recovered within the ensembles, thereby providing confidence in our sampling strategy.

First, we evaluated whether PROTAC sampling (Step 2) could generate crystal-like conformations. Using both bound and unbound states, the MC sampling produced PROTAC conformations with free warhead RMSD values below our 2 Å threshold (**Figure 3A, Table S1**), confirming compatibility with crystal-like PPD poses. Additionally, we observed that the initial PROTAC conformation from Step 1 influenced TC retrieval, emphasizing the need to sample from multiple PROTAC clusters for optimal results (**Figure S3**).

**Figure 3.**
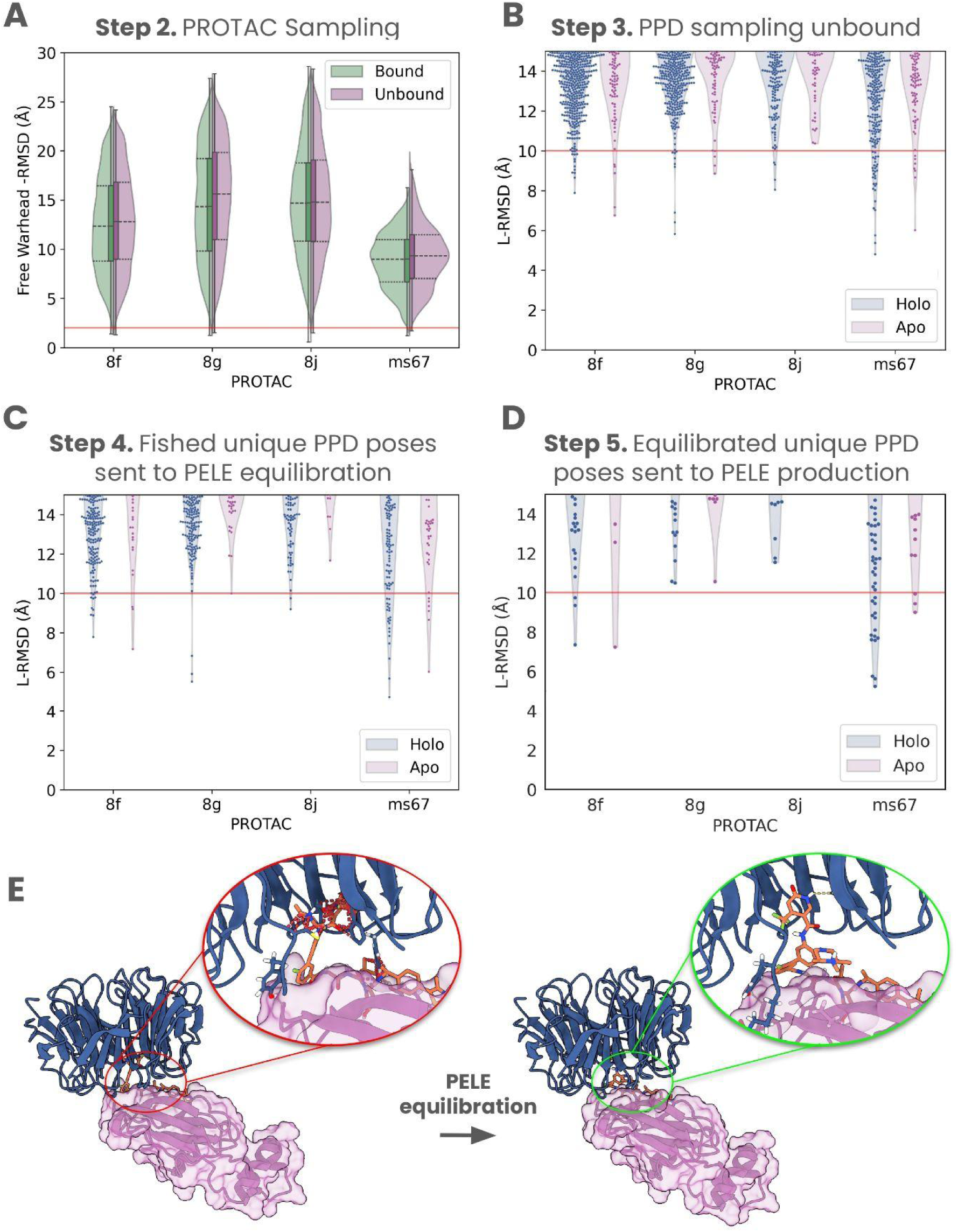
Tracking NN poses at different filtering stages of the protocol using unbound protein conformations. **(A)** Distribution of free warhead RMSD values compared to each PROTAC crystal structure. Conformations with RMSD below 2 Å (red line) are considered compatible with crystal-like PPD poses. **(B)** L-RMSD distribution of the PPD poses obtained by FTDock with holo and apo modes (see **Figure S4** for the bound case). **(C)** L-RMSD distribution of unique fished PPD poses using unbound crystals (see **Figure S12** for the bound case). **(D)** L-RMSD distribution of the unique PPD poses selected for PELE production (see **Figure S13** for the bound case). The red lines in the previous panels indicate the 10 Å cut-off distance used to define NN poses. **(E)** Fished TC structure with clashes shown in red dashed lines (left), and same crystal after PELE equilibration simulation (right), where the free warhead is accommodated, forming hydrogen bonds (yellow dashed lines).

In parallel, we examined whether the PPD (Step 3) could sample near native (NN) poses, defined as ligand-protein root mean square distance (L-RMSD) below 10 Å relative to the TC crystal structure, as in previous studies.^22,23^ Using both holo and apo PPD sampling modalities, we assessed bound (**Figure S4**) and unbound (**Figure 3B**) protein conformations (see **Table S2** for RMSD comparison). In all cases except for 8j in the apo unbound mode, FTDock successfully retrieved NN conformations (**Figure 3B**, **Table 1**). This suggests that the apo mode is more likely to fail when PROTAC-induced TCs involve few protein-protein contacts and there is no pre-existing interaction between the proteins (**Figure S5**). While this challenge also exists in the holo mode, sampling poses with different PROTAC conformations yields some NN solutions, aligning with the biophysical process where TC formation follows binary complex formation.

**Table 1:** Number of unique NN and PPD poses at different filtering stages for unbound conformations. **Refer to Table S3** for results using bound conformations. Abbreviations: Equi - PELE Equilibration, Prod. - PELE Production.

| PROTAC<br>(NN / Unique<br>PPD poses) | 8f |  | 8g |  | 8j |  | ms67 |  |
| --- | --- | --- | --- | --- | --- | --- | --- | --- |
|  | Holo | Apo | Holo | Apo | Holo | Apo | Holo | Apo |
| **Step3**<br>**PPD** | 15 /<br>554,400 | 5 /<br>92,400 | 7 /<br>554,400 | 4 /<br>92,400 | 6 /<br>554,400 | 0 /<br>92,400 | 28 /<br>369,600 | 7 /<br>92,400 |
| **Step3**<br>**Filtered** | 15 /<br>61,588 | 5 /<br>8,906 | 7 /<br>56,909 | 4 /<br>8,906 | 6 /<br>45,806 | 0 /<br>8,906 | 28 /<br>39,461 | 7 /<br>8,906 |
| **Step 4**<br>**Fished** | 9 /<br>5,195 | 3 /<br>876 | 3 /<br>5,354 | 1 /<br>1,082 | 2 /<br>5,170 | 0 /<br>1,318 | 20 /<br>1,242 | 6 /<br>415 |
| **Step 5**<br>**Equi.** | 208 /<br>5,195 | 17 /<br>876 | 244 /<br>5,354 | 13 /<br>1,082 | 79 /<br>5,170 | 10 /<br>1,318 | 54 /<br>1,242 | 7 /<br>415 |
| **Step 6**<br>**Prod.** | 25 /<br>367 | 1 /<br>37 | 41 /<br>749 | 7 /<br>74 | 5 /<br>200 | 2 /<br>40 | 29 /<br>374 | 3 /<br>77 |

At this stage, we also assessed whether pyDock^38^ interaction energy—which includes desolvation, Coulombic electrostatics, and van der Waals energy between the ligand protein and the receptor protein (including the PROTAC in holo mode)^39^ —could be used to identify NN poses. The results revealed that, although FTDock sampling retrieved NN poses in most cases, only few had stabilizing negative interaction energy. In holo mode, NN poses generally have positive energy values, making energy-based selection challenging (**Figure S6**). This results from van der Waals energy penalties caused by PROTAC-induced clashes at the interface, which a rigid body PPD software cannot release, leading to highly energetic conformations. Moreover, the lack of flexibility in the PPD algorithm makes the sampling highly dependent on the PROTAC initial conformation (**Figure S7**). Therefore, extensive sampling from multiple initial PROTAC conformations was crucial to ensure NN pose identification. However, while holo mode increased the absolute number of NN poses compared to apo mode, the overall NN percentage was sometimes reduced by up to 3.75-fold (**Table 1, Table S3**).

To shortlist our TCs candidates, we applied a two-step filtering process to eliminate PPD poses incompatible with TC formation. This filtering assessed whether BS were approximately face-to-face and within a feasible distance for PROTAC bridging (**Figure S1**). We applied a uniform BS distance threshold across all PROTACs, as previous studies found TC formation to be independent of linker length.^40^ Despite being relatively tolerant, these filters significantly increased the percentage of NN poses by up to 12-fold respect the original PPD data (**Table 1**, **Figure S8**) without excluding any NN pose (**Table 1**, **Figure S9**).

Further filtering PPD poses with PROTAC sampling conformations (Step 4) substantially enriched the NN ratio by up to 27-fold compared to the filtered PPD poses pool and up to 224-fold respect to the original PPD sampling (**Table 1**, **Figure S10**). However, this selection criteria for identifying potential TC candidates resulted in the loss of some NN poses (**Figure 3C**), in part due to subtle deviations in the free warhead’s spatial alignment, exceeding the 2 Å RMSD threshold.

To resolve steric clashes in our fished TCs, we applied semi-flexible MC equilibrations with PELE (Step 5), which successfully eliminated clashes in the selected TCs (**Figure 3E**). This local refinement significantly increased the number of NN poses by up to 40-fold. Notably, for 8j compound simulation in apo mode with unbound conformation, equilibration recovered NN poses from an initial PPD set that originally contained none (**Table 1**), suggesting that these conformations heavily depend on the presence of the PROTAC. These results highlight the effectiveness of our equilibration protocol, mostly based in coupling the sampling of the PROTAC dihedral space with the side chain neighbour rotamer libraries (see Methods), in guiding the system toward local minima, including crystal-like conformations, by simultaneously modeling both the free protein and the PROTAC.

Although the number of unique NN poses increased after equilibration, not all were advanced to production (**Figure 3D**). This is because our selection criteria prioritized the lowest energy poses with the free warhead positioned near its BS. Keep in mind that we are not exclusively interested in reproducing NN or crystallographic poses, but rather an energy landscape exploration of potential TCs. Despite this reduction in the number of NN poses to start the MC production stage (Step 6), all the PROTACs had at least a simulation converging to a crystal-like structure (**Table 1**), further demonstrating the capability of our MC protocol to reach local minima consistent with experimental data.

In conclusion, our results highlight the robustness of our workflow in generating a broad range of potential TC complexes, including NN ones (**Figure S14**). This in depth exploration of TCs landscape could provide structural insights to aid in the rational optimization of PROTACs.

### Correlation of TC stability based on TC frequency

Having established the ability of our workflow to generate NN poses while exploring the structural landscape of TCs, we next investigated whether filtering and counting the number of TCs per PROTAC could align with its thermodynamic TC stability, as similarly proposed by previous studies to correlate with PROTAC degradation capacity.^23,24^

Strictly speaking, the ternary K_D_ can be derived by the ratio of TC occurrences relative to all other possible microstates, weighted by the Boltzmann factor. We therefore hypothesized that the number of TCs in our TC landscapes could align with the experimental ternary K_D_, as it captures the collective contribution of all possible conformations in the landscape. Nevertheless, exhaustively sampling the entire conformational space is not feasible in practice. Therefore, our workflow aims to implement an important sampling approach on the TC landscape, focusing on a specific subspace of conformations that could potentially lead to TC formation. As a result, our method might be better suited to congeneric series of PROTACs, where non-TC conformational spaces could be more similar.

In our first analysis, we counted potential TCs generated using a purely geometric approach (Step 4) and found a correlation between the number of TCs and linker length, with the longest PROTACs, 8g and 8j, which yield the highest numbers (**Table 2, Table S4**). These results indicate that simpler geometric approaches that do not account for more in-depth biophysical simulations may induce length-correlated artifacts. To escape from this source of error, our analysis was focused only on those TCs produced through PELE equilibration (Step 5) and production (Step 6). The results then aligned more closely with the ternary K_D_. Notably, the shortest PROTAC, ms67, emerged as the PROTAC generating the highest number of TCs followed by 8g and 8f across all bound/unbound and apo/holo study cases. In contrast, 8j consistently yielded fewer TCs than 8f, the least TC-stabilizing PROTAC, which might be indicating that the landscapes are not comparable due to the different nature of 8j’s linker (aromatic) respect to the other aliphatic linkers.

**Table 2:** Number of TC candidates and number of unique PPD poses that form a TC candidate at different filtering stages for unbound conformations. PROTACs are ordered from high to low ternary K_D_ (see Figure 2). Refer to **Table S4** for results using bound crystals.

| PROTAC<br>(Total TCs /<br>Unique PPD poses) | 8f |  | 8g |  | 8j |  | ms67 |  |
| --- | --- | --- | --- | --- | --- | --- | --- | --- |
|  | Holo | Apo | Holo | Apo | Holo | Apo | Holo | Apo |
| **Step 4**<br>**Fished** | 62,959 /<br>5,195 | 7,336 /<br>876 | 65,389 /<br>5,354 | 8,950 /<br>1,082 | 66,215 /<br>5,170 | 10,818 /<br>1,318 | 28,312 /<br>1,242 | 7,984 /<br>415 |
| **Step 5**<br>**Equi.** | 94 /<br>31 | 1 /<br>1 | 280 /<br>65 | 43 /<br>9 | 13/<br>6 | 0 | 2,565 /<br>192 | 891 /<br>52 |
| **Step 6**<br>**Prod.** | 3,019 /<br>54 | 68 /<br>4 | 7,128 /<br>137 | 1,883 /<br>15 | 293 /<br>16 | 68 /<br>4 | 13,444 /<br>131 | 5,642/<br>37 |

Altogether, the results suggest that filtering TCs with energy-based criteria leads to better correlations with ternary K_D_. Simply counting TCs assumes equal contribution, whereas their actual influence depends on energy via the Boltzmann factor. Considering only energetically feasible TCs makes this approximation more accurate. Additionally, the dynamic nature of TCs seen experimentally supports this assumption, as it implicitly implies similar energy levels. Nonetheless, this counting approach seems to be sensitive to the nature of the linker, as exemplified by 8j, where, the rigidity of the aromatic linker may significantly alter the system’s energy landscape, thus, complicating comparisons with the other more flexible linkers. Next, we moved beyond simply counting TCs to examine whether the energy landscapes of these TCs provide deeper insights into TC stability.

### Correlation of TC stability based on TC energy landscapes

To investigate TC energy landscapes, we analyzed the energy profiles obtained during the PELE production phase (Step 6). The initial TC candidates used for the simulations enabled extensive sampling of the landscapes, covering conformations with L-RMSD values ranging from ∼5 to ∼40 Å relative to their crystal structures (**Figures S15 to S22**). Interestingly, PROTACs 8g and ms67 (to a greater extent) exhibited a strong ability to form TCs across a broad range of conformations while maintaining similar total energy values (**Figures S19 to S22**), suggesting that these PROTACs induce highly dynamic TCs. In contrast, 8j primarily formed TCs at specific conformations (L-RMSD ∼10 Å and ∼30 Å), likely due to the rigidity of its linker.

One could hypothesize that a PROTAC capable of accommodating a wider range of TC conformations might facilitate faster TC formation. However, experimental data on TC formation (**Figure 2**), suggest that conformational flexibility alone is insufficient since 8j and ms67 have similar values. In reality, to establish a direct correlation, we would need to simulate the TC formation process itself. Instead, our approach primarily samples the conformational space of potential final TCs, omitting TC formation from the binary complex. Yet, capturing these initial processes would require significantly more extensive sampling.

Once a TC is formed, the focus shifts to its thermodynamic stability. We observed that, within our conformational subspace, largely biased toward potential TC poses, these energy landscapes tend to converge into approximately 2 to 10 distinct minimas, representing distinct PROTAC-induced TC conformations (**Figure 4B,D,E-K**, **Figure S23**). Notably, ms67 (**Figure 4L**) and 8g consistently form NN TCs, whereas 8j and 8f exhibit greater difficulty in doing so. Nevertheless, crystal-like conformations did not always lead to the lowest energy minimas within the explored landscape (**Figure 4B,D,E**), as also observed in previous studies.^32^ This underscores the importance of exploring alternative binding modes beyond crystals to fully understand PROTAC mediated TCs.

**Figure 4.**
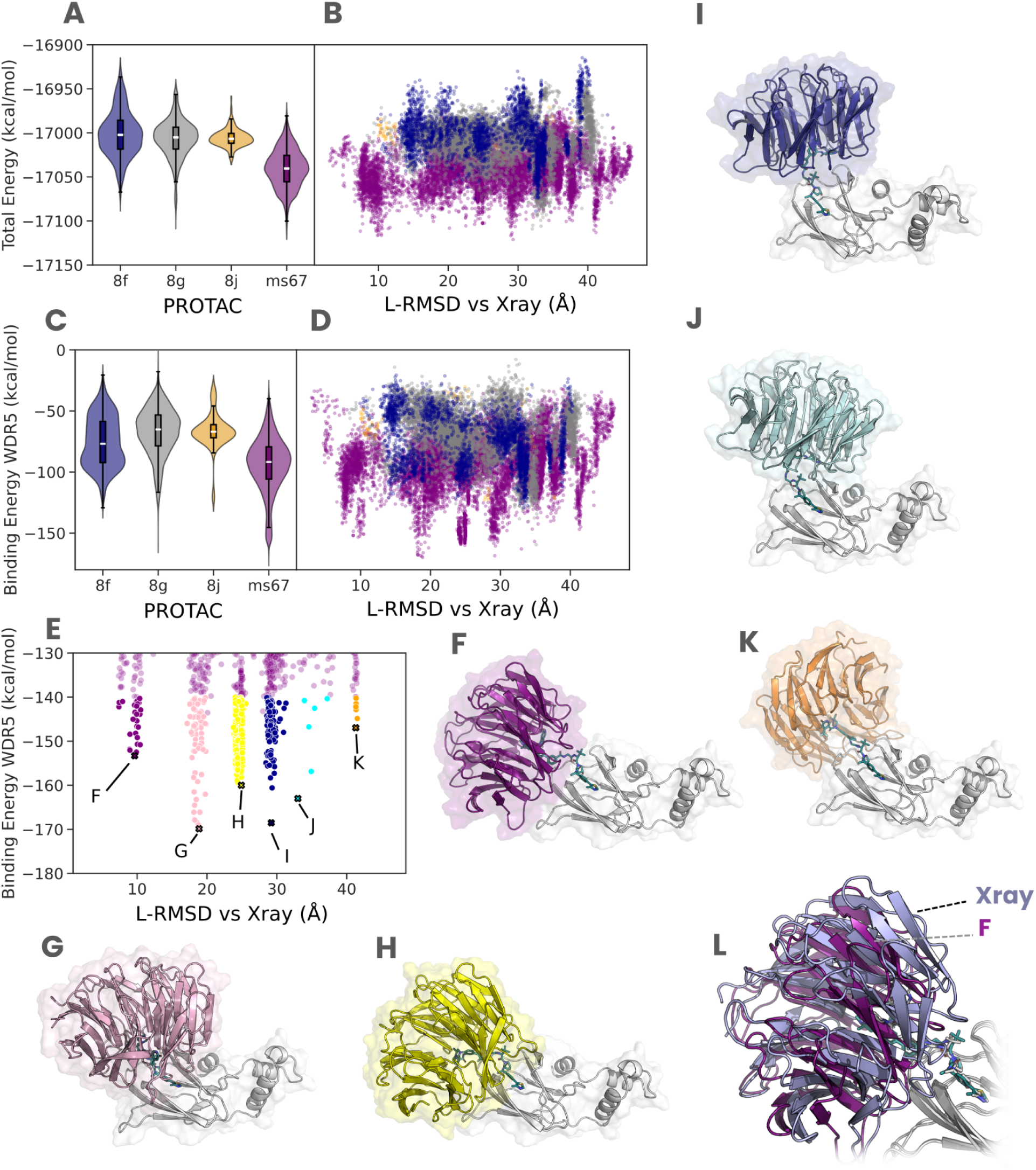
TC energy landscapes for unbound conformation using holo mode. **(A,B)** Total energy and **(C,D)** binding energy profiles for each PROTAC. The violin plots of **(A,C)** represent the energy values distribution shown in **(B,D)**. **(E)** Zoom in ms67 binding energy landscape to show low energy minima. **(F-K)** Lowest energy binding modes for ms67. **(L)** Model **(F)** aligned on top of VHL of the X-ray 7JTP (WDR5 in light blue and VHL in gray).

The energy distribution of the landscapes clearly distinguishes ms67 as the PROTAC that induces the most stable TCs (**Figure 4A,C**, **Figure S24**), consistent with the experimentally observed TC half-life (**Figure 2**). This suggests that once a TC forms, better binding energies indicate higher likelihood of remaining stable, thus leading to higher TC half-life values, as seen with ms67. However, ms67’s enhanced stability in both ternary K_D_ and TC half-life may be due to multiple factors: it not only forms the lowest-energy TCs, but it is also compatible with a broader range of TC conformations, potentially reducing entropic penalties.

The TC landscape analysis provided insights into both the energetic and dynamic nature of PROTAC-mediated TCs. Energetically, it revealed that PROTACs play a crucial role in stabilizing TCs, reinforcing their importance in promoting target proximity. From a dynamic perspective, the TC energy landscapes appear to capture the TC half-life once formed, suggesting its dependence on binding affinity of the TC. Thus, the energy distribution within these landscapes complements TC frequency-based methods for evaluating PROTAC stability from a kinetic perspective, further aiding in the rational design and optimization of PROTACs.

## Conclusions

In this work, we presented a PPD-MC-based workflow for the comprehensive exploration of the TC landscape. Our results demonstrate that these landscapes can be leveraged to: i) identify energetically relevant TC conformations, including the crystal. ii) compare thermodynamic TC stability described by the ternary K_D_ obtained by similar PROTACs, and iii) estimate kinetic stability of the TC described by TC half-life. This information can help reduce experimental workload in PROTAC optimization, which is largely reliant on trial-and-error approaches.^18^

Nevertheless, our method faces challenges in recovering NN poses when protein contacts are limited, particularly in the apo mode. This suggests that TCs entirely dependent on PROTAC bridging (i.e., where no direct protein-protein interactions exist) are difficult to detect using workflows that rely on PPD for TC candidates generation. Additionally, the nature of the PROTAC linker constrains the approach’s applicability in predicting thermodynamic TC stability. This is because our approach estimates the ternary K_D_ based on the number of relevant TCs but omits other non-TC possible microstates, which are implicitly captured by experimental K_D_ and can be significantly affected by linker properties. Furthermore, even with MC techniques, exploring TC landscapes remains CPU intensive, limiting its use to screening small PROTAC series with tens of compounds.

Overall, considering both the strengths and limitations of our approach, it is best suited for optimizing short PROTACs’ TC stability, particularly those where protein-protein interactions play a role. This aligns with the industry’s focus on developing drug-like PROTACs with favorable properties.

## Methods

### Dataset

Four PROTACs from Schwalm et al.^10^ were selected as the primary systems to benchmark our computational workflow. These PROTACs utilize VHL E3 ligase to promote degradation of WDR5 protein. They were selected due to their comprehensive experimental data, including ternary and binary dissociation constants, degradation metrics, and available crystal structures (**Figure 2**). Interestingly, despite their chemical similarity, this subset spans a wide range of ternary K_D_ values, making it ideal for testing the performance of our computational approach on PROTACs prioritization task based on TC formation.

### System preparation

Protein structures were obtained from the Protein Data Bank (PDB). Specifically, we used 7JTP for the bound VHL-WDR5 system and 5NVX and 4QL1 for the unbound VHL and WDR5, respectively. These unbound structures were chosen because their proteins were co-crystallized with the used warheads and had been used in previous studies for unbound-state analysis.^32,41^ All protein and protein/PROTAC systems were prepared with the Protein Preparation Wizard from Schrödinger^42^ to fill missing side-chains and protonate the system at pH 7 with PROPKA.^43^

### PROTAC docking using the least buried warhead

PROTAC docking is performed with Glide from Schrödinger^44^ with an inner box size of 20 Å and outer box of 35 Å. Given that binding modes are often well-known, we use that information to set core constraints and attach the warhead to the receptor and to focus the conformational sampling to the linker and outer warhead region. In total, for each PROTAC we generate up to 20 different conformations. We used the volume overlap clustering tool from Schrödinger to group redundant conformations and get a diverse set of binary complex conformations. PROTAC poses were linked using the centroid and the optimal number of clusters was measured based on the Kelley penalty.^45^

### PROTAC conformational sampling using PELE

For each binary complex, we independently sample the PROTAC conformational space using PELE side-chain perturbation scheme. In this protocol, each PELE step applies random rotations in three rotamers of the linker and free warhead, while keeping the bound warhead rotamers fixed. This is followed by a side-chain prediction step to release clashes between the PROTAC and the receptor protein. Finally, the step is accepted or rejected following the Metropolis criterion. In this study, we performed 31 independent trajectories for each initial binary complex, with each trajectory consisting of 200 PELE steps. This comprehensive sampling process requires approximately one hour of computational time.

### PPD with FTDock

For holo PPD, the PROTACs are parametrized with the general AMBER force field (GAFF)^46^ using ANTECHAMBER 22 through AmberTools.^47^ Then, for both holo and apo modes, we prepare the inputs with pyDock^38^ and run FTDock^29^ to sample the conformational space, thereby generating 92400 poses. To compute the interaction energies for all FTDock generated poses we use pyDock scoring function.

### Filter PPD poses by binding site orientation and distance

To filter PPD poses, we first define the BS plane for both the target protein and the E3 ligase by manually selecting three residues on each protein. The warhead exit vector is approximated as the normal vector to this plane (**Figure S1A,B)**. To ensure the correct orientation of the exit vector (ie., pointing toward the solvent), we manually select a fourth residue located within the protein core to ensure the exit vector points outward.

For each PPD pose, we compute the dot product between the BS normal vectors of the E3 ligase and the target protein. Poses with a dot product greater than 0 (indicating the vectors point in the same direction) are discarded (**Figure S1C**). We then compute the distance between the centers of the BS planes (**Figure S1D**), defined as the geometric centers of the alpha carbons of the three selected residues defining the BS plane. Only poses with BS plane centers separated by 30 Å or less are retained (**Figure S1E**). This ensures proper orientation, proximity of BSs, and preservation of pose diversity.

To optimize computational efficiency and minimize data generation, we apply rotation and translation matrices directly, avoiding the need to generate PDB files for each pose.

### Generation of TCs candidates

Assuming that the warhead binding modes are well-known, we superpose the receptor protein warhead of all PROTACs conformations to the PPD poses. Then, we compute a three atom RMSD of the free protein warhead with respect to its known binding mode for each PROTAC conformation and PPD pose pair. The three atoms are manually selected and given as input to the workflow (**Figure S2**). Our recommendation is selecting atoms that are representative for the whole warhead conformation and not for a single rigid region (e.g., atoms in an aromatic ring that are in the same plane). The selection of spread atoms in the warhead reduces the number of TC candidates and prevents conformations with excessive clashes. We consider that a TC candidate is obtained when we find a PPD pose whose BSs can be linked with a PROTAC conformation with free warhead RMSD < 2 Å. From this pool of TCs, for each unique PPD pose, we selected the TC with the lowest free warhead RMSD to be sent to the PELE equilibration step. The poses selected for the equilibration step contain the PPD pose with the PROTAC in the conformation that resulted in the lowest free warhead RMSD.

### TC equilibration with PELE

We use PELE to release steric clashes and obtain low-energy states from the previously generated TCs candidates. Each equilibration is performed using 12 independent trajectories. Each trajectory consists of 10 PELE steps, with random translations (0.3 Å and 0.1 Å) and rotations (0.01 and 0.03 rads) applied to the free protein, along with random rotations (maximum angle: 30° per rotatable bond) of the linker and free warhead rotamers. After each PELE perturbation, interface side-chains are optimized followed by a minimization step to relieve newly introduced clashes in the perturbation step. Importantly, no distance constraints are applied to bias the simulations toward TC formation. During the simulation, we monitor two specific distances between the free warhead and its BS (e.g., potential hydrogen bond interactions, see **Figure S25**). These distances are used to validate the formation of TCs, based on a predefined distance threshold. Each simulation requires approximately 1 hour of computation and 12 CPUs per initial TC candidate.

### Selection of equilibrated TCs

For each unique PPD pose, we select the equilibrated TC candidate with the lowest total energy. A potential TC candidate is considered eligible if the two monitored distances between the free warhead and its BS are below 12 Å. Among these, only the top 50% with the lowest total energy are advanced to the production step.

### Exploration of TC energy landscape

For each equilibrated TC we perform a PELE simulation with 63 independent trajectories, each consisting of 10 *AdaptivePELE* epochs and 8 PELE steps per epoch. During each PELE step, random translations (0.3 Å and 0.1 Å) and rotations (0.01 and 0.03 radians) are applied to the free protein, along with random rotations of the linker, this time without a maximum rotation angle to enable unrestricted sampling. As in the equilibration step, no distance constraints are used.

To enhance sampling toward TCs mediated by the PROTAC, all sampled conformations are clustered by the PROTAC contact map at the end of each epoch. Representative structures for the next epoch are selected to minimize a specific distance between the free warhead and its BS (preferably a distance tracking a buried interaction). This is achieved by setting the *epsilon* parameter in *AdaptivePELE* to 90%, which ensures that 90% of the selected poses minimize the distance, while the remaining 10% explore less frequent low-density clusters. This balance allows the system to sample both optimal and less explored configurations.

### Analysis of TC energy landscapes

Conformations are considered to form TCs when the two distances between free warheads and its BS are below 5 Å (**Figure S25**). Data from PELE simulations was analyzed using Pandas.^48^ TC energy landscapes were plotted using the structures which pass the threshold, using seaborn^49^ and matplotlib.^50^ Additionally, structures were observed and illustrated using ChimeraX^51^ or PyMOL.^52^

## ASSOCIATED CONTENT

The following files are available free of charge. Supplementary Figures and Tables (PDF)

The workflow code to prepare and run pyDock, FTDock and PELE jobs will be available in the following GitHub repository upon paper acceptance: https://github.com/annadiarov/PELETAC

## AUTHOR INFORMATION

### Author Contributions

The manuscript was written through contributions of all authors. All authors have given approval to the final version of the manuscript.

### Notes

The authors declare the following competing financial interest(s): At the time the work described in this manuscript was carried out, C.P-L., J.M-P., C.P., L.D. and V.G. were employees of Nostrum Biodiscovery. The remaining authors report no competing interests.

## ACKNOWLEDGMENT

We would like to thank Víctor Montal from Barcelona Supercomputing Center and Ilia Lecha, former Nostrum Biodiscovery employee, for interesting discussions. We also thank the Spanish Ministry of Science and Innovation for the predoctoral fellowship grant to A.M.D-R (FPU/03921).

## ABBREVIATIONS

BS: Binding Site
K_D_: dissociation constant
L-RMSD: Ligand protein Root Mean Square Distance
MD: Molecular Dynamics
MC: Monte Carlo
MM/GBSA: molecular mechanics with generalized Born and surface area solvation
NN: Near Native
PDB: Protein Data Bank
PELE: Protein Energy Landscape Exploration
POI: Protein Of Interest
PPD: Protein-Protein Docking
PROTAC: Proteolysis-Targeting Chimeras
TC: Ternary Complex
TPD: Target Protein Degradation

