## Supplementary Information for "Characterization of PROTACs by Ternary Complex Landscape Exploration using Monte Carlo simulations"

### Supplementary Figures

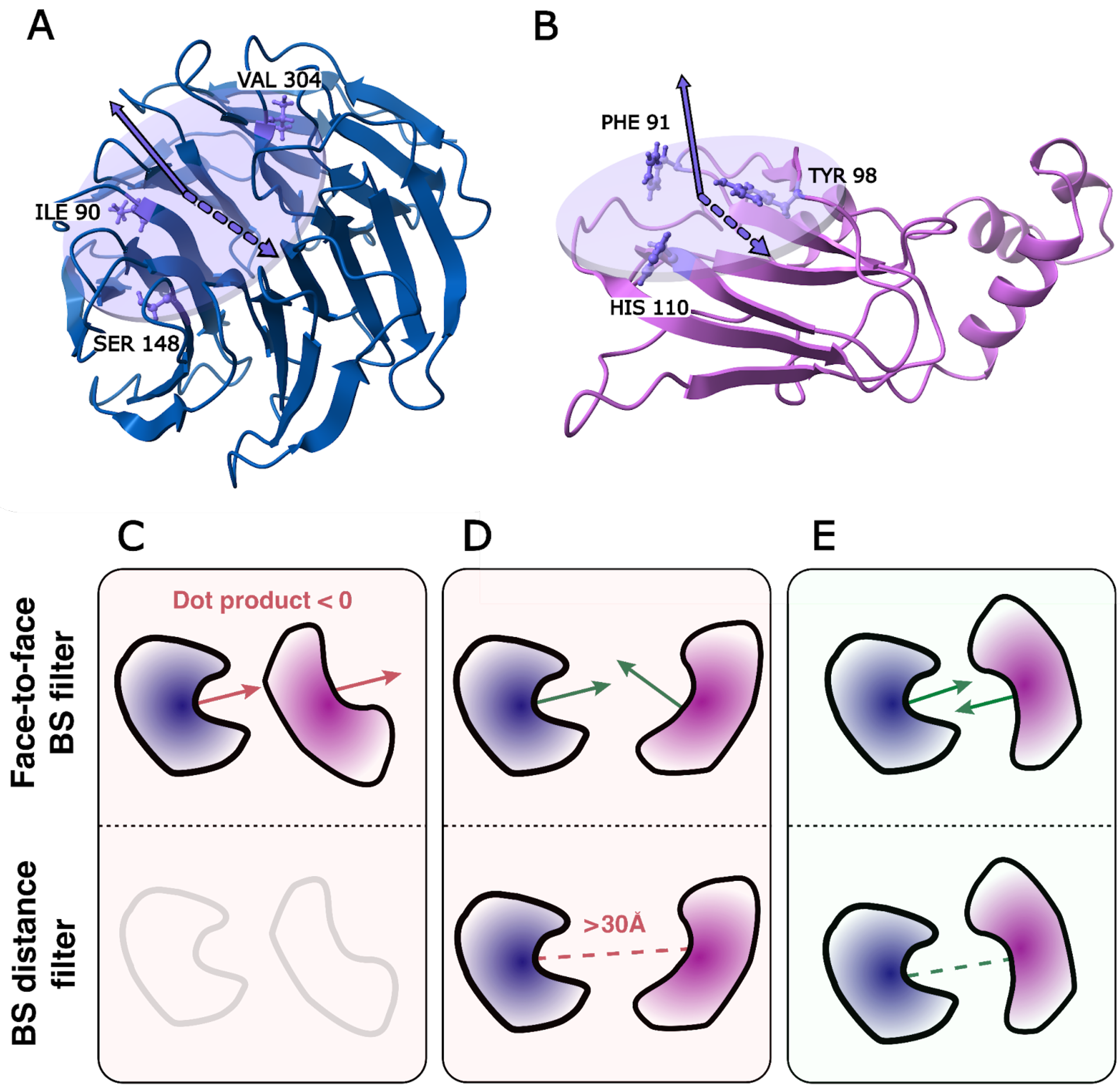


**Figure S1. PPD filtering by BS and distance orientation.** Schematic representation showing that the BS vector is perpendicular to the BS plane in **(A)** WDR5 and **(B)** VHL respectively. The carbon alpha atoms from the residues shown were used to define the BS plane. The filtering of PPD poses involves the following steps: **(C)** first discarding PPD poses where the warhead BSs are not face-to-face (dot product > 0), and **(D)** second removing poses where the BS plane center is farther than 30 Å. **(E)** Finally, poses satisfying the BS and distance orientation filters are kept.

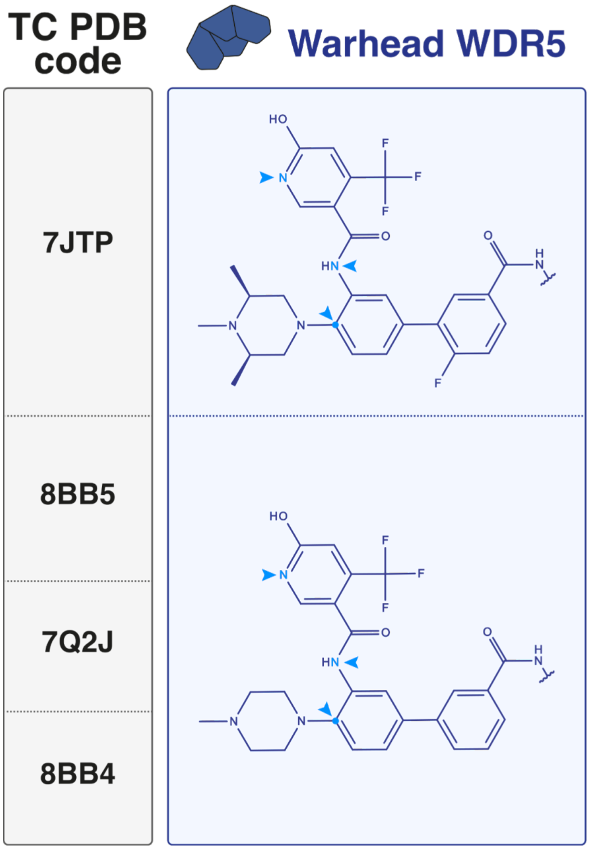


**Figure S2. Atoms used to compute free warhead RMSD.** The arrows indicate the WDR5 selected atoms.

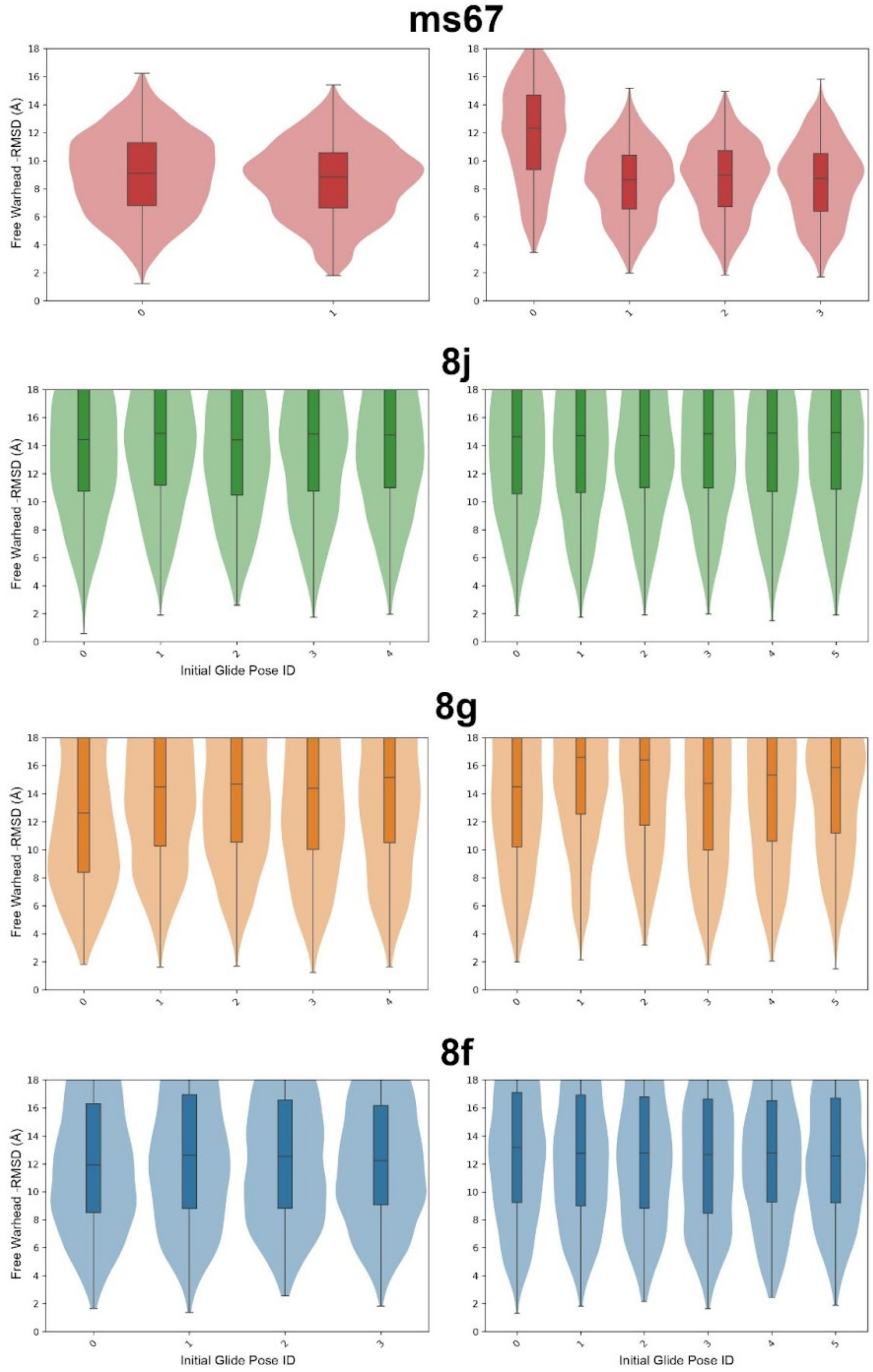


**Figure S3. Three point free warhead RMSD distributions of PROTAC sampling conformations relative to the crystal structure.** The PROTAC is docked at the VHL binding site, leaving the WDR5 warhead free. Each PROTAC shows as many RMSD distributions as its number of initial Glide docked poses. Distributions on the left correspond to the bound VHL conformation, while those on the right refer to the unbound conformation.

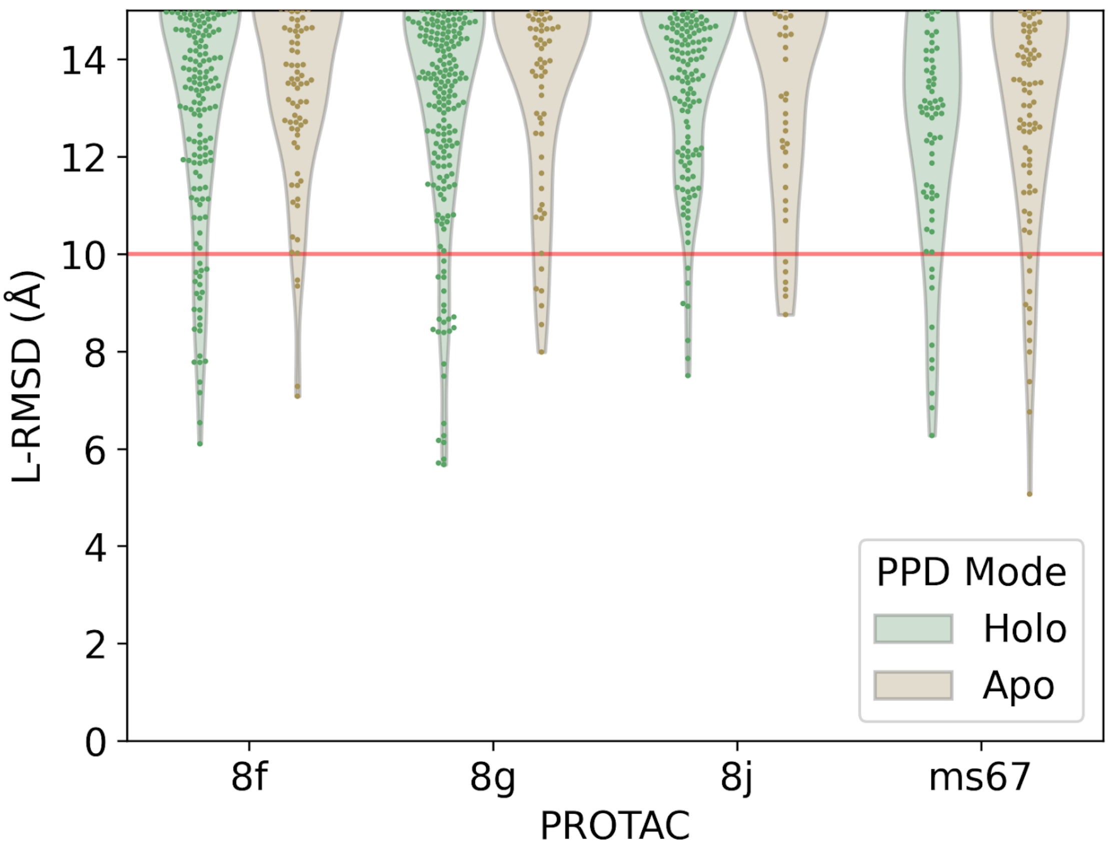


**Figure S4. PPD L-RMSD distribution using bound conformations.** Each point represents a PPD obtained by FTDock, Notice that the holo mode distributions contain all PPD poses obtained with all initial PROTAC clusters. For the distribution obtained using unbound crystals, refer to **Figure 3B**.

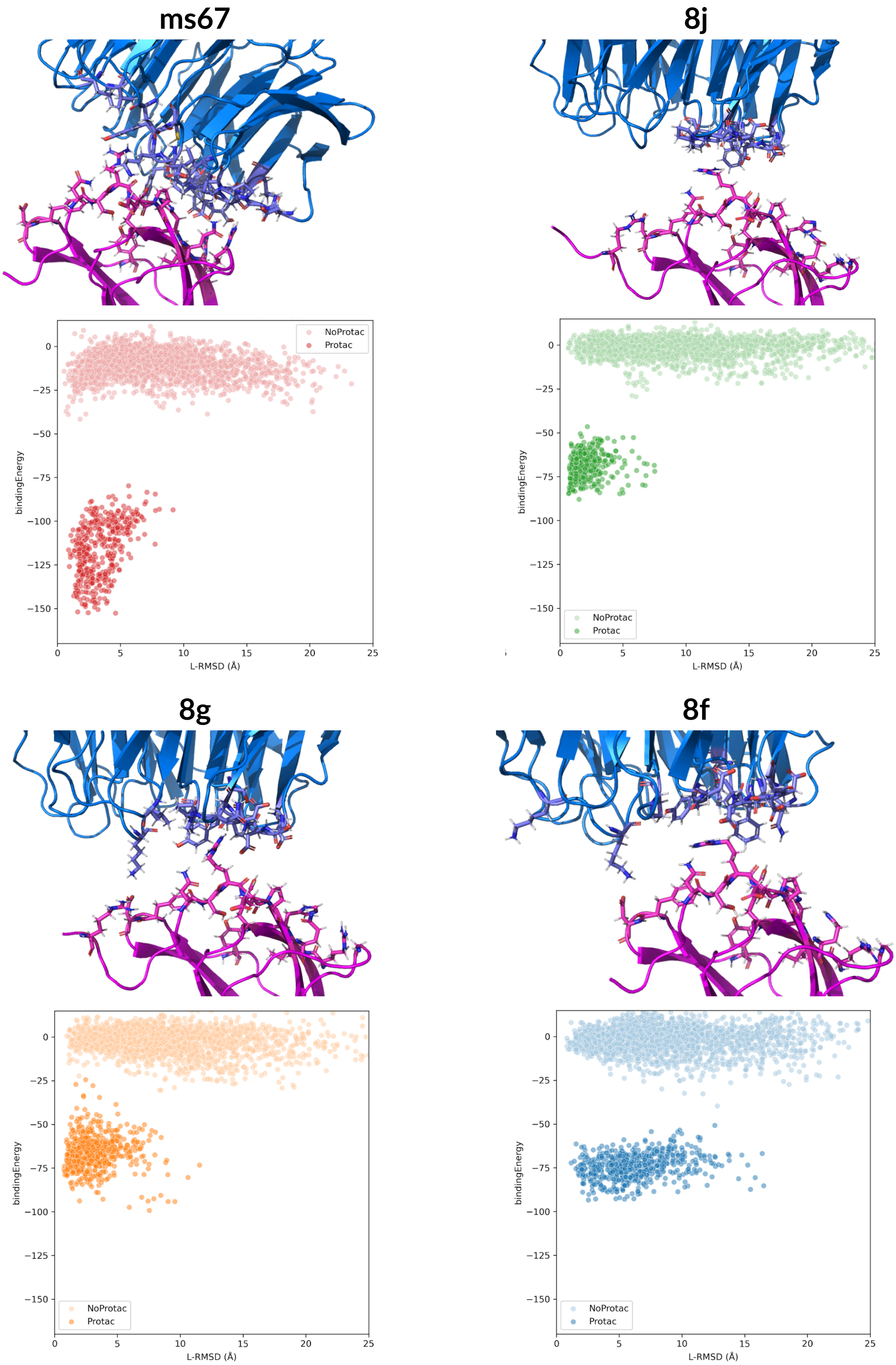


**Figure S5: Characterization of VHL-WDR5 crystals.** Crystals interfaces for each PROTAC are shown on top. VHL is displayed in pink, while WDR5 in blue. Interface residues are shown as sticks. PELE local refinement energy profiles for the crystals with and without PROTAC are shown below.

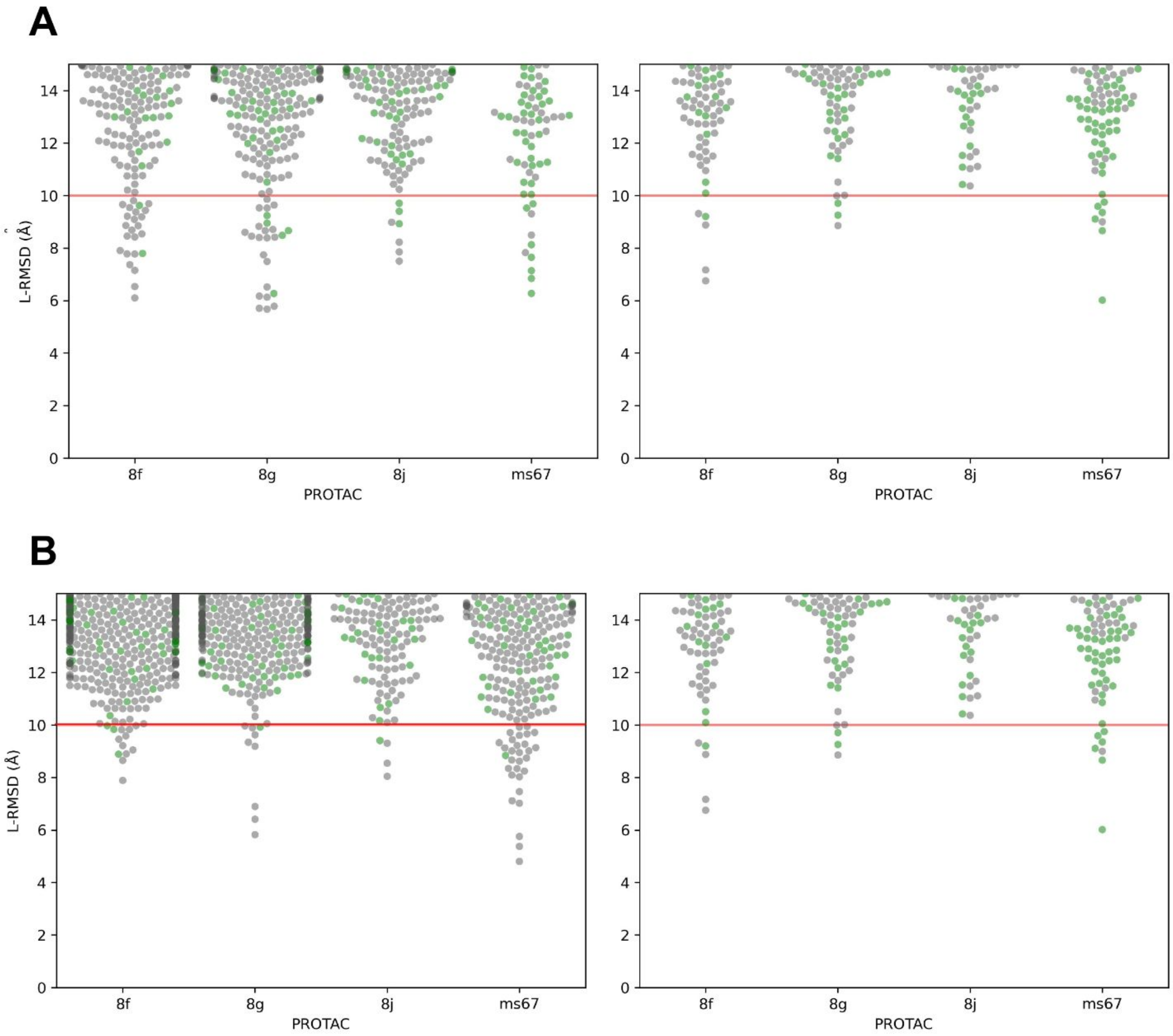


**Figure S6. L-RMSD PPD distributions, with green indicating poses with negative pyDock interaction energy. (A)** Distributions using the bound conformation in holo (left) and apo (right) modes. **(B)** Distributions using the unbound conformation in holo (left) and apo (right) modes.

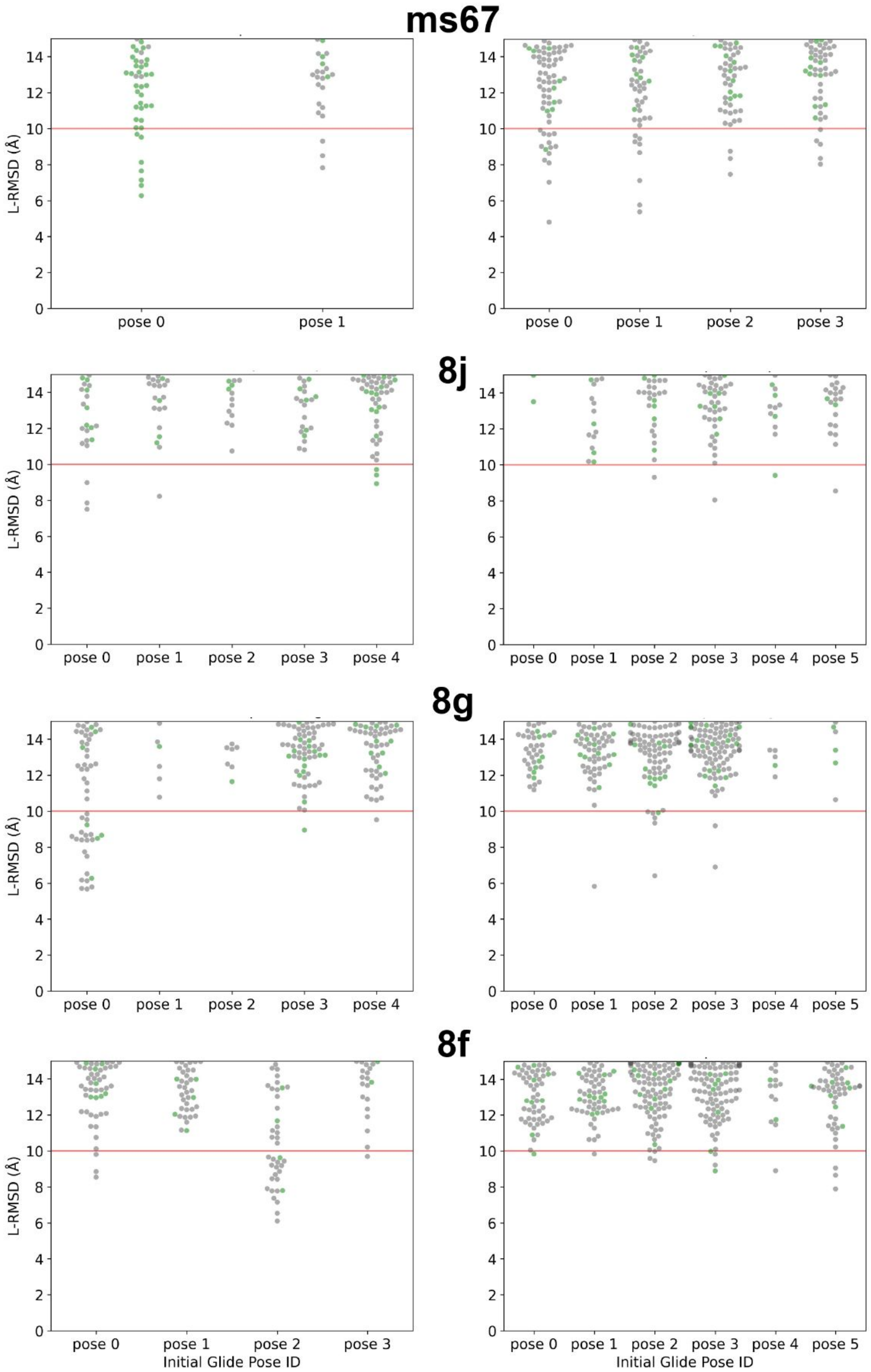


**Figure S7. L-RMSD distributions of holo PPD, with green indicating poses with negative pyDock interaction energy.** Each PROTAC has multiple distributions based on its initial Glide-docked poses. Left: bound conformations. Right: unbound conformations.

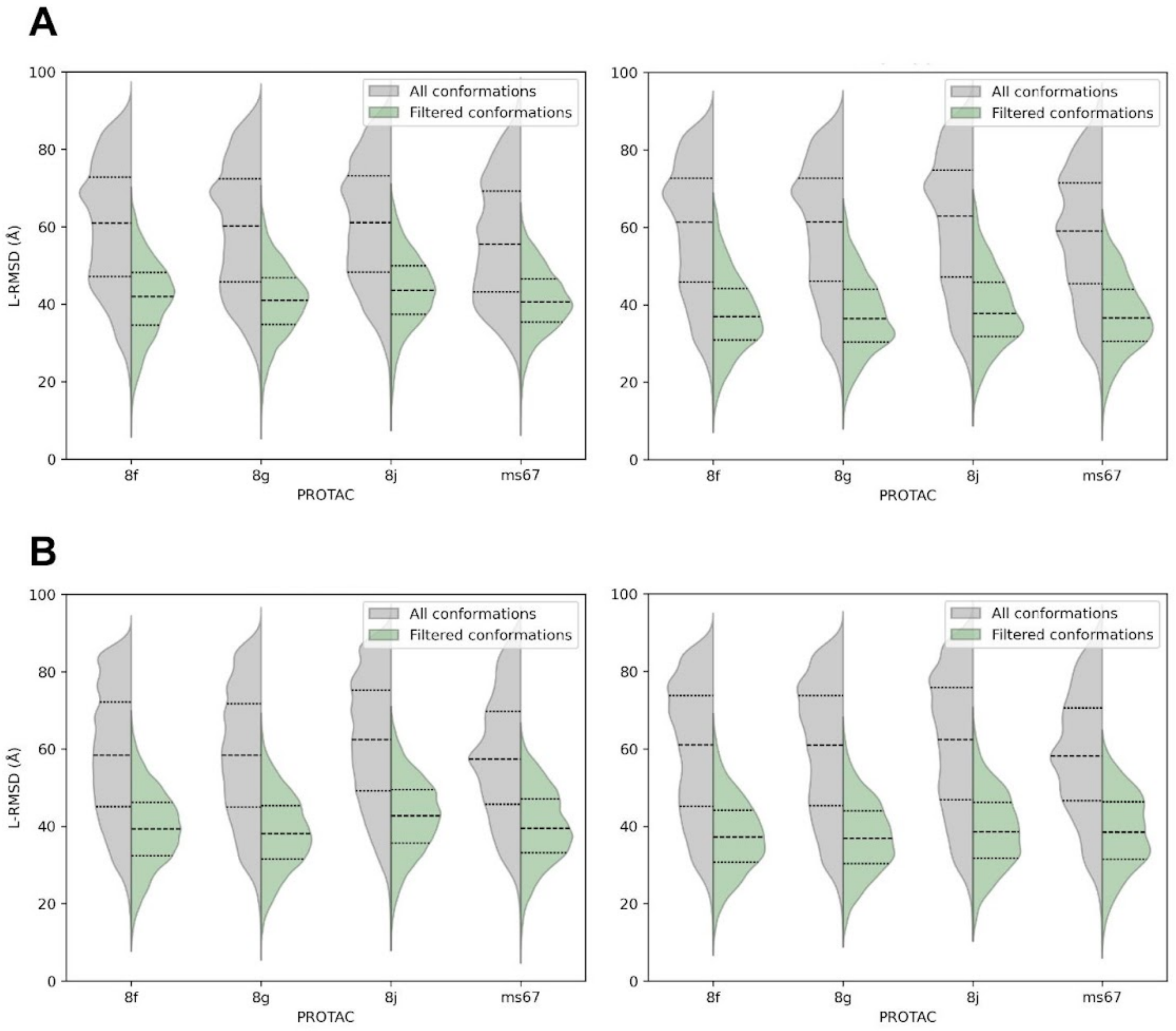


**Figure S8. Comparison of L-RMSD distributions for poses that pass versus those that fail the binding site orientation and distance filters.** **(A)** Distributions based on the bound conformation in holo (left) and apo (right) modes. **(B)** Distributions based on the unbound conformation in holo (left) and apo (right) modes.

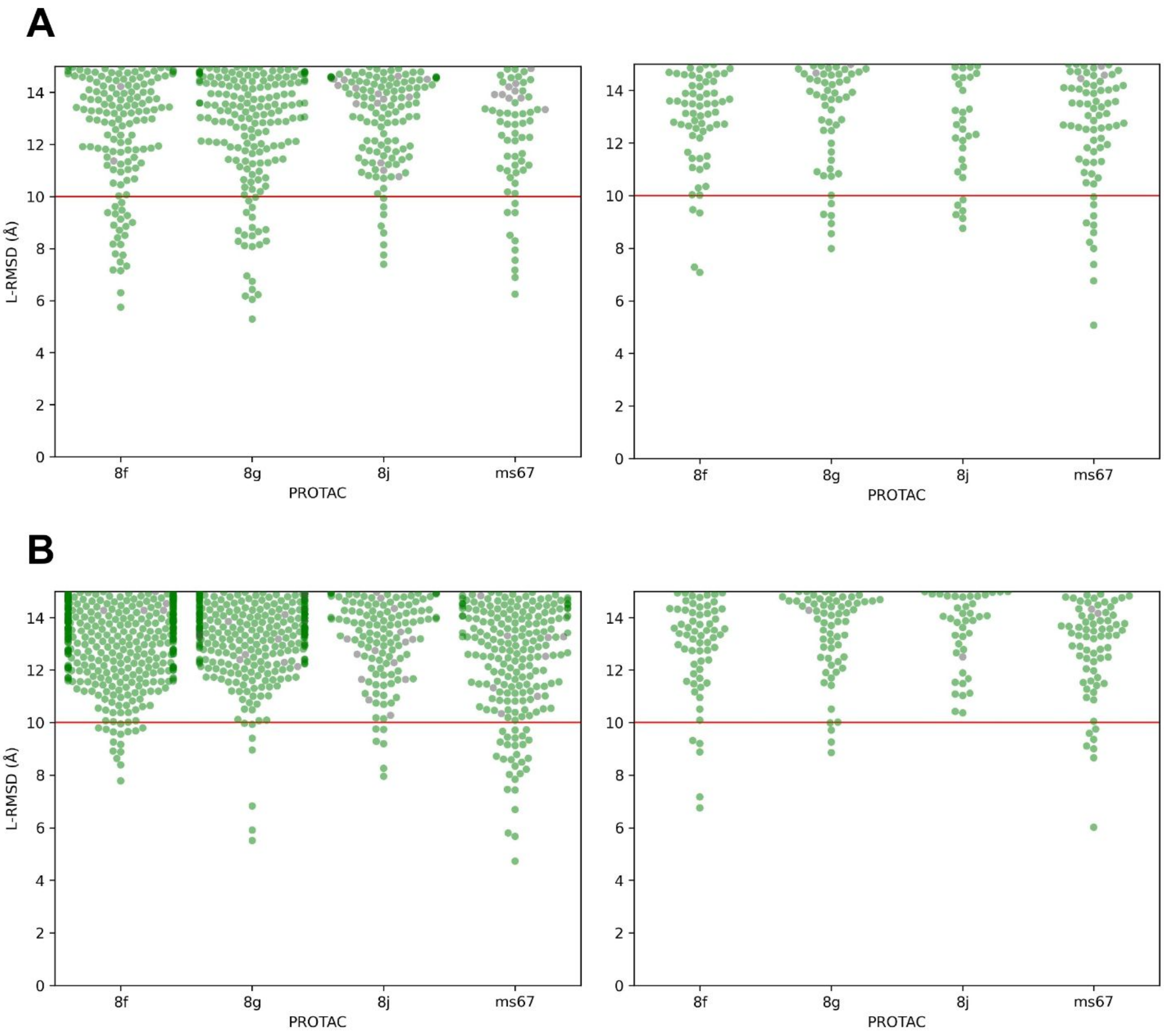


**Figure S9. L-RMSD PPD distributions, with green indicating poses that pass the binding site orientation and distance filters. (A)** Distributions using the bound conformation in holo (left) and apo (right) modes. **(B)** Distributions using the unbound conformation in holo (left) and apo (right) modes.

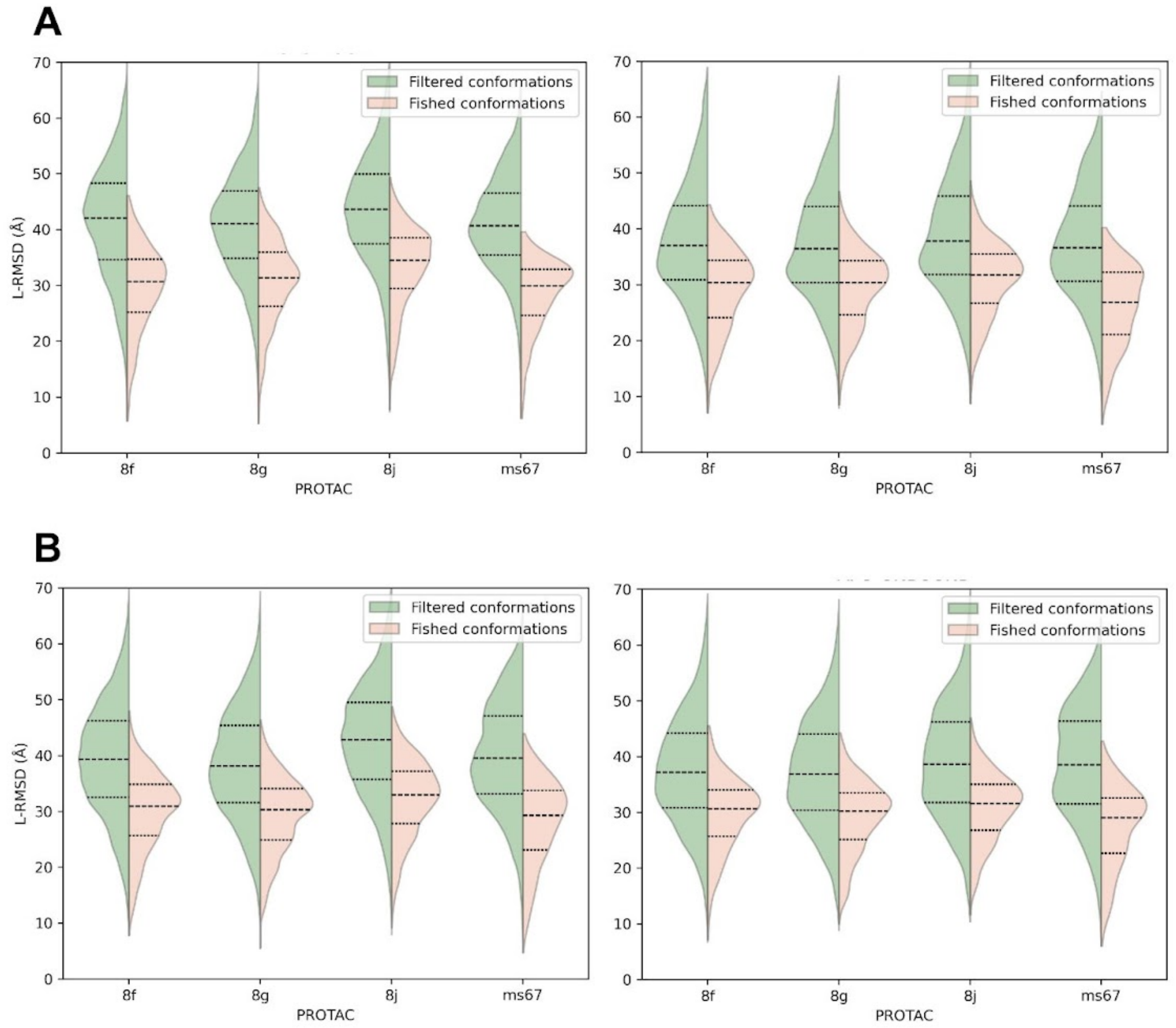


**Figure S10. Comparison of L-RMSD distributions between fished and filtered poses.** **(A)** L-RMSD distributions using bound conformation in holo (left) and apo (right) modes. **(B)** L-RMSD distribution obtained from PPD with unbound conformation in holo (left) and apo (right) modes.

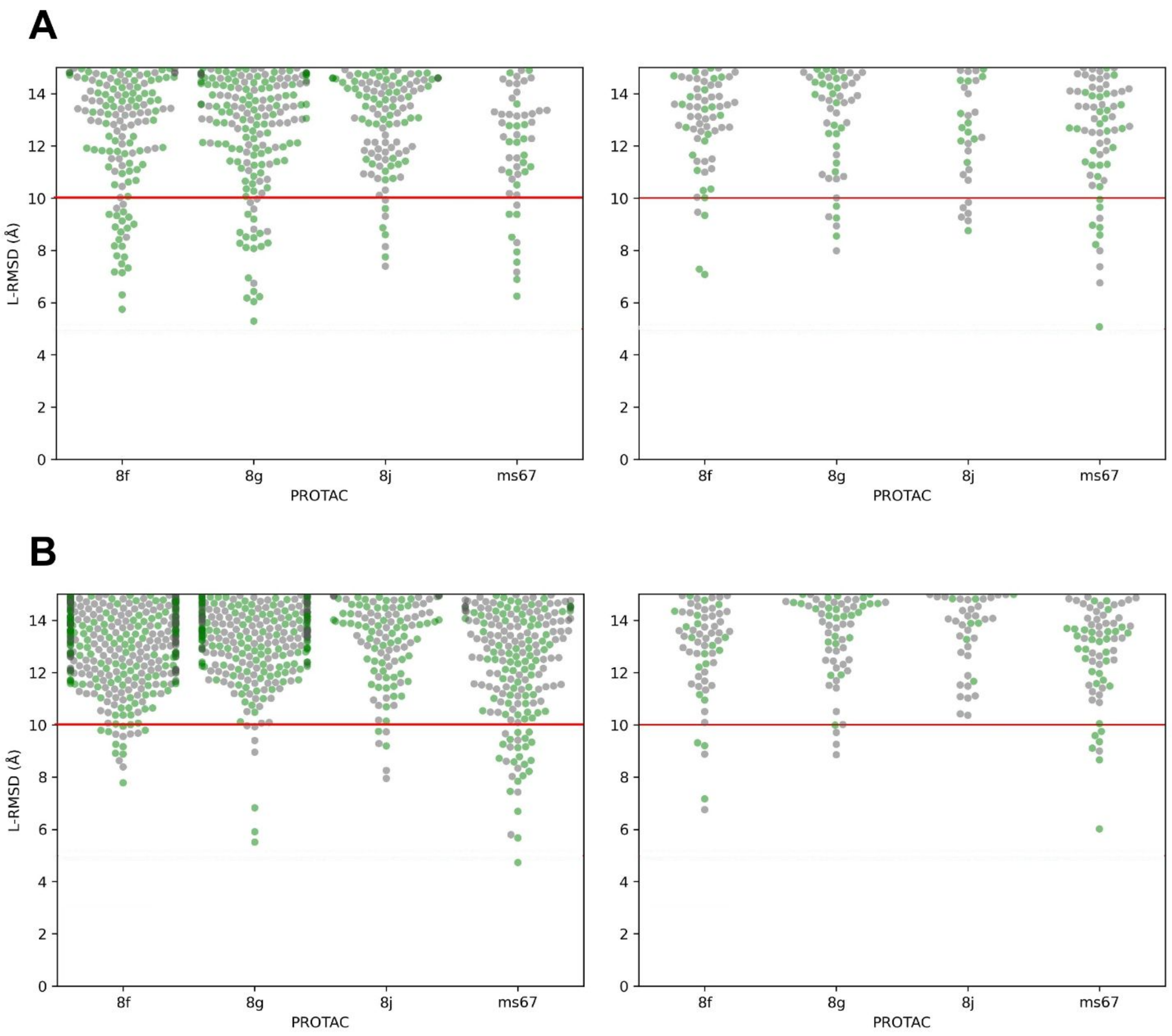


**Figure S11. L-RMSD PPD distributions colored based on compatibility with at least one PROTAC sampling conformation.** **(A)** L-RMSD distributions using the bound conformation in holo (left) and apo (right) modes. **(B)** L-RMSD distributions using the unbound conformation in holo (left) and apo (right) modes. Green dots represent PPD poses compatible with at least one PROTAC sampling conformation, while gray dots indicate poses that passed the orientation filters but were not compatible with any PROTAC conformation.

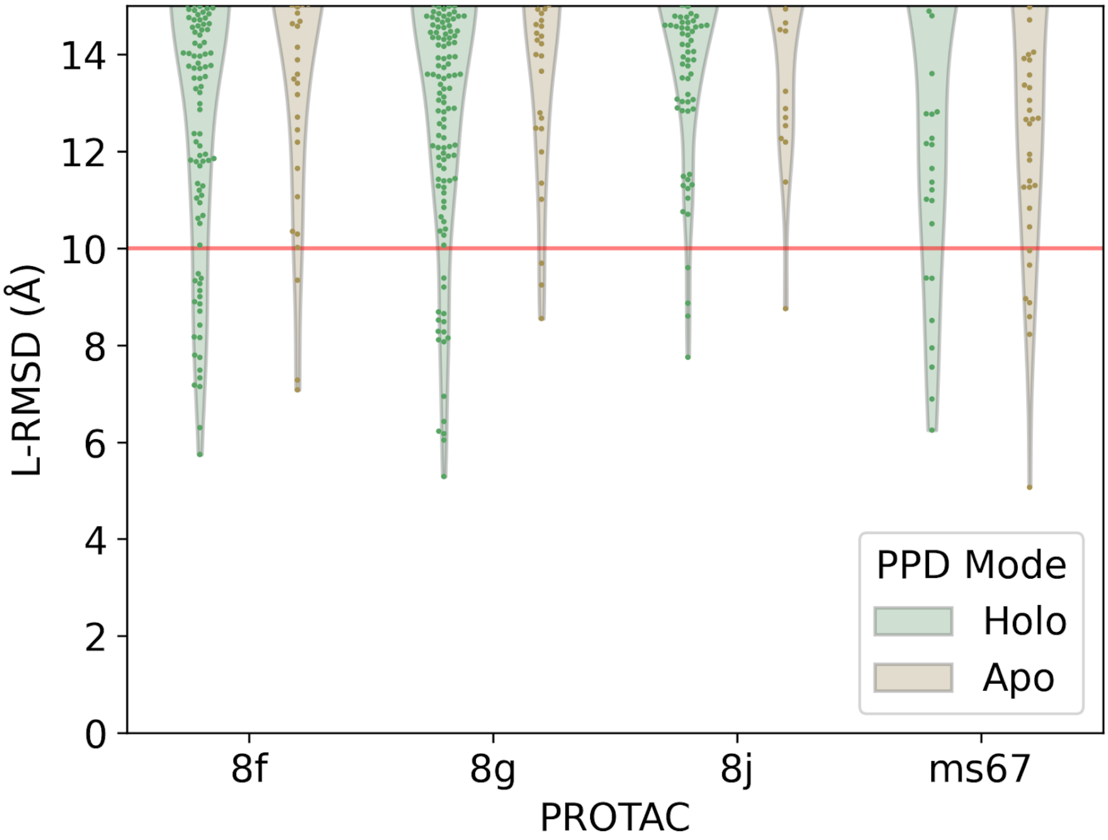


**Figure S12. L-RMSD distribution of fished PPD poses when using bound conformations.** Each point represents a PPD pose that was compatible with at least one conformation from the PROTAC sampling step. For the distribution obtained using unbound crystals, refer to **Figure 3C**.

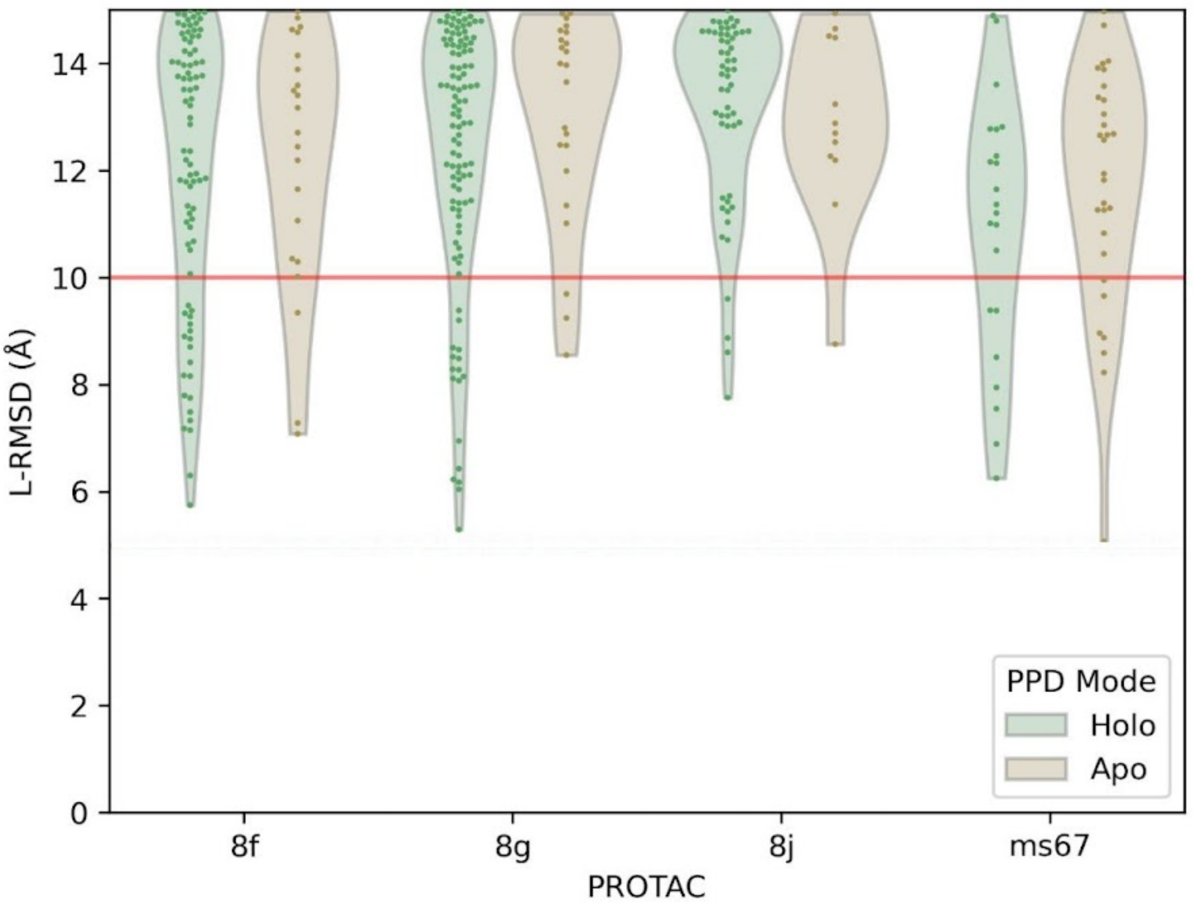


**Figure S13. L-RMSD distribution of unique fished PPD poses selected for PELE production when using bound conformations.** Each point represents a fished PPD pose that passed the filtering criterion after the PELE equilibration. For the distribution obtained using unbound protein conformation, refer to **Figure 3C**.

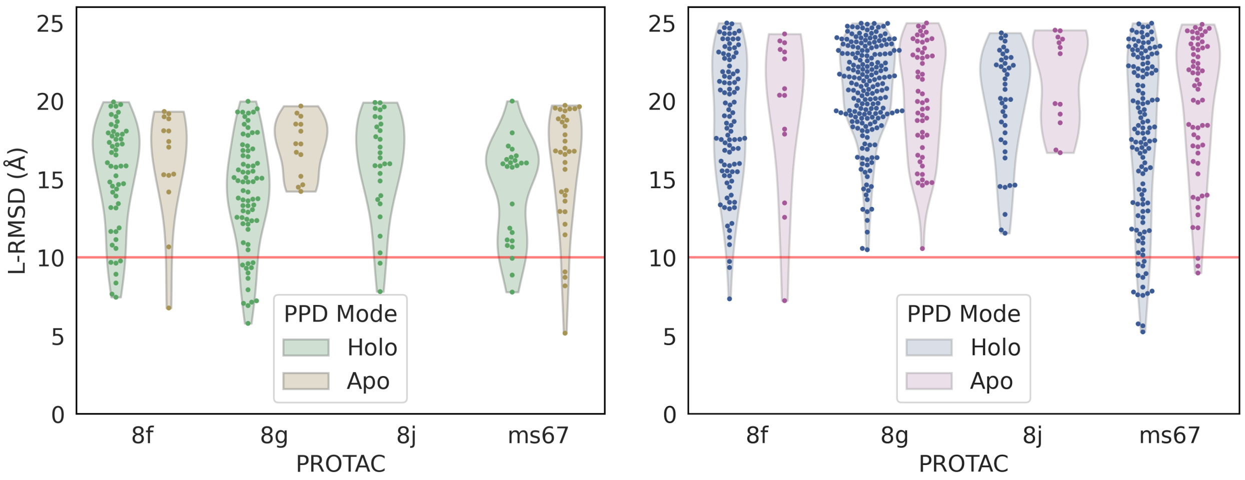


**Figure S14. All unique PPD poses selected for PELE production.** Each point represents a fished PPD that passed the filtering criterion after the PELE equilibration for the bound (left) and unbound (right) protein conformations.

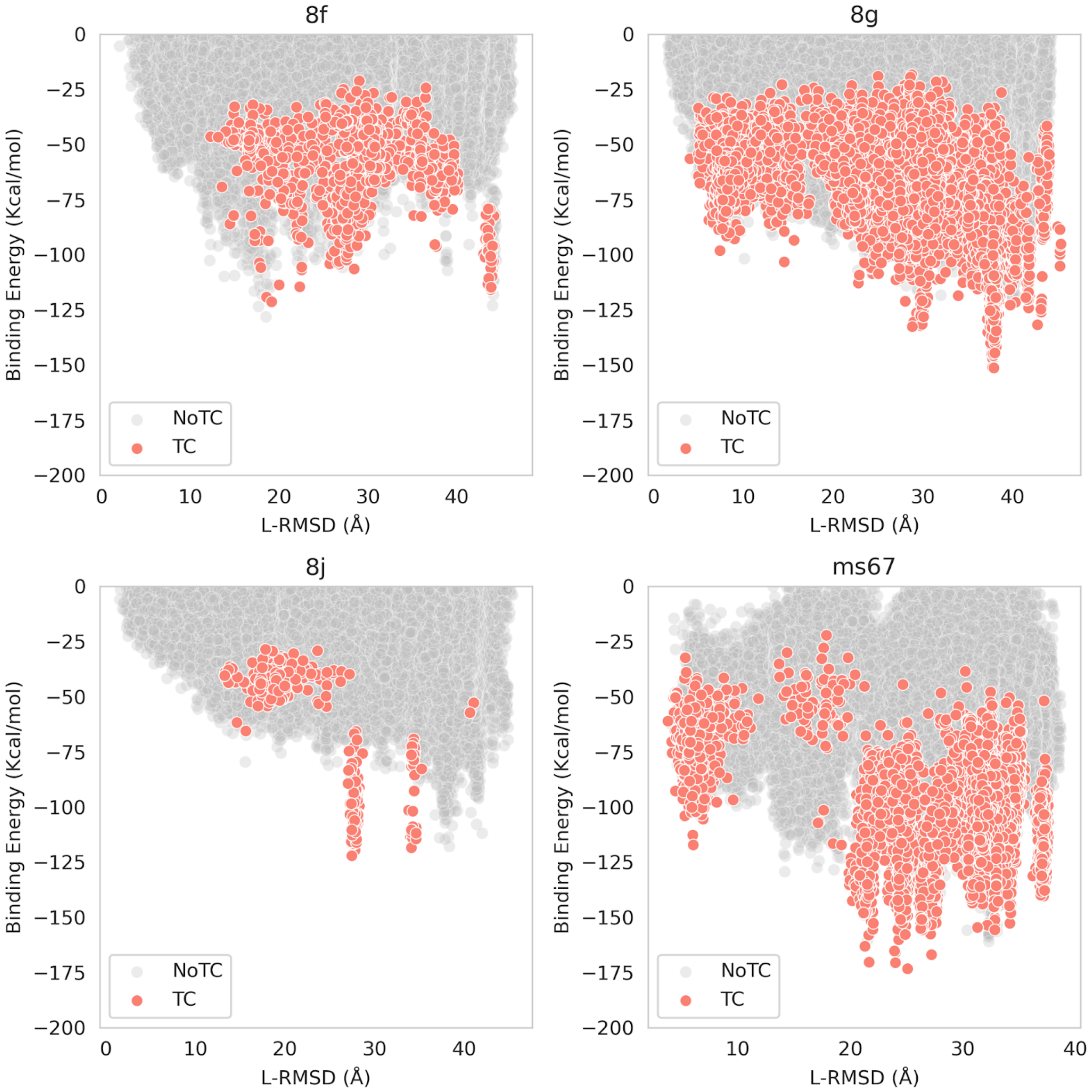


**Figure S15: Holo bound binding energy TC landscapes**. Each point represents a system conformation accepted by PELE that forms a TC (orange) or not (gray).

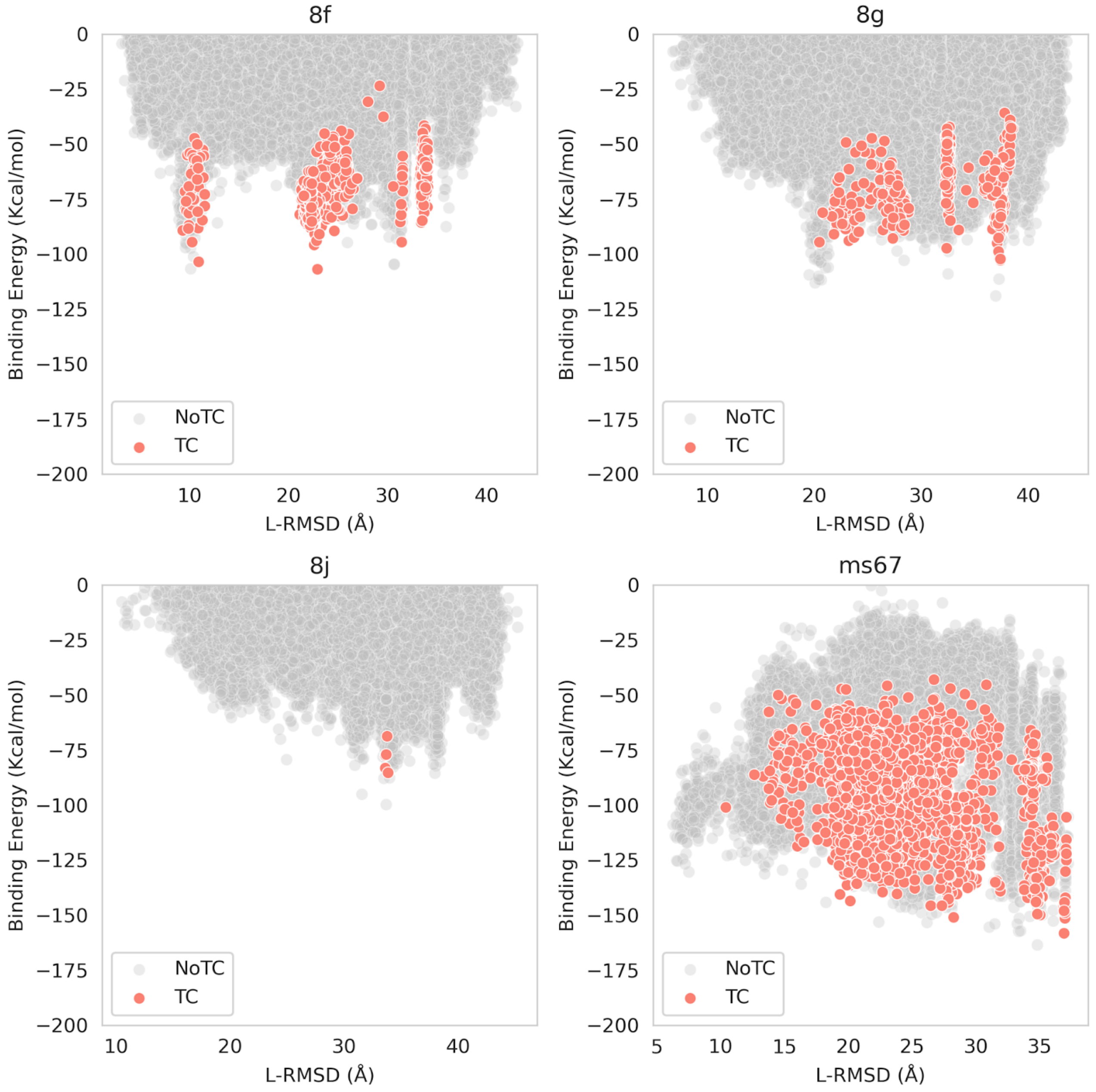


**Figure S16: Apo bound binding energy TC landscapes**. Each point represents a system conformation accepted by PELE that forms a TC (orange) or not (gray).

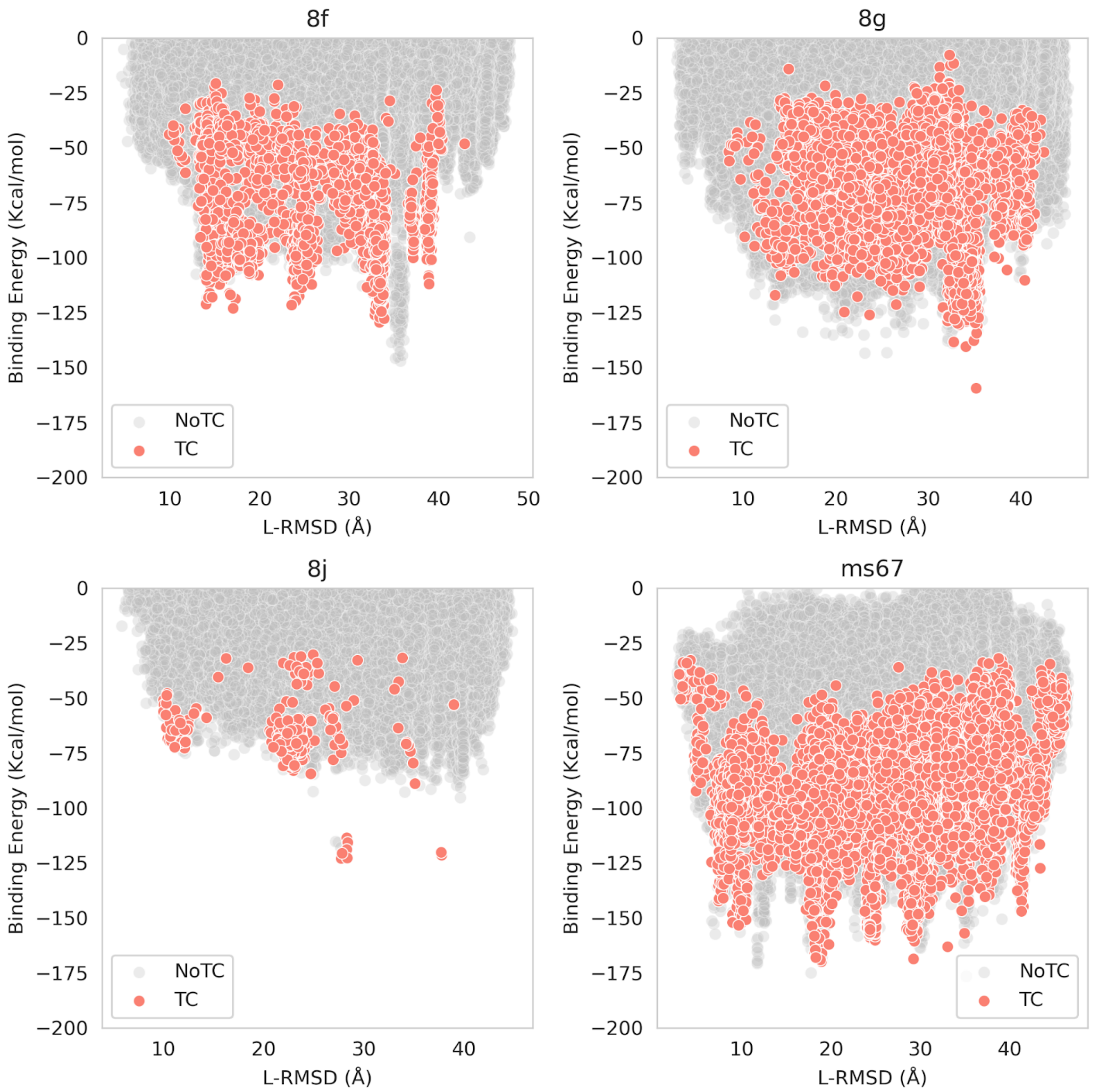


**Figure S17: Holo unbound binding energy TC landscapes**. Each point represents a system conformation accepted by PELE that forms a TC (orange) or not (gray).

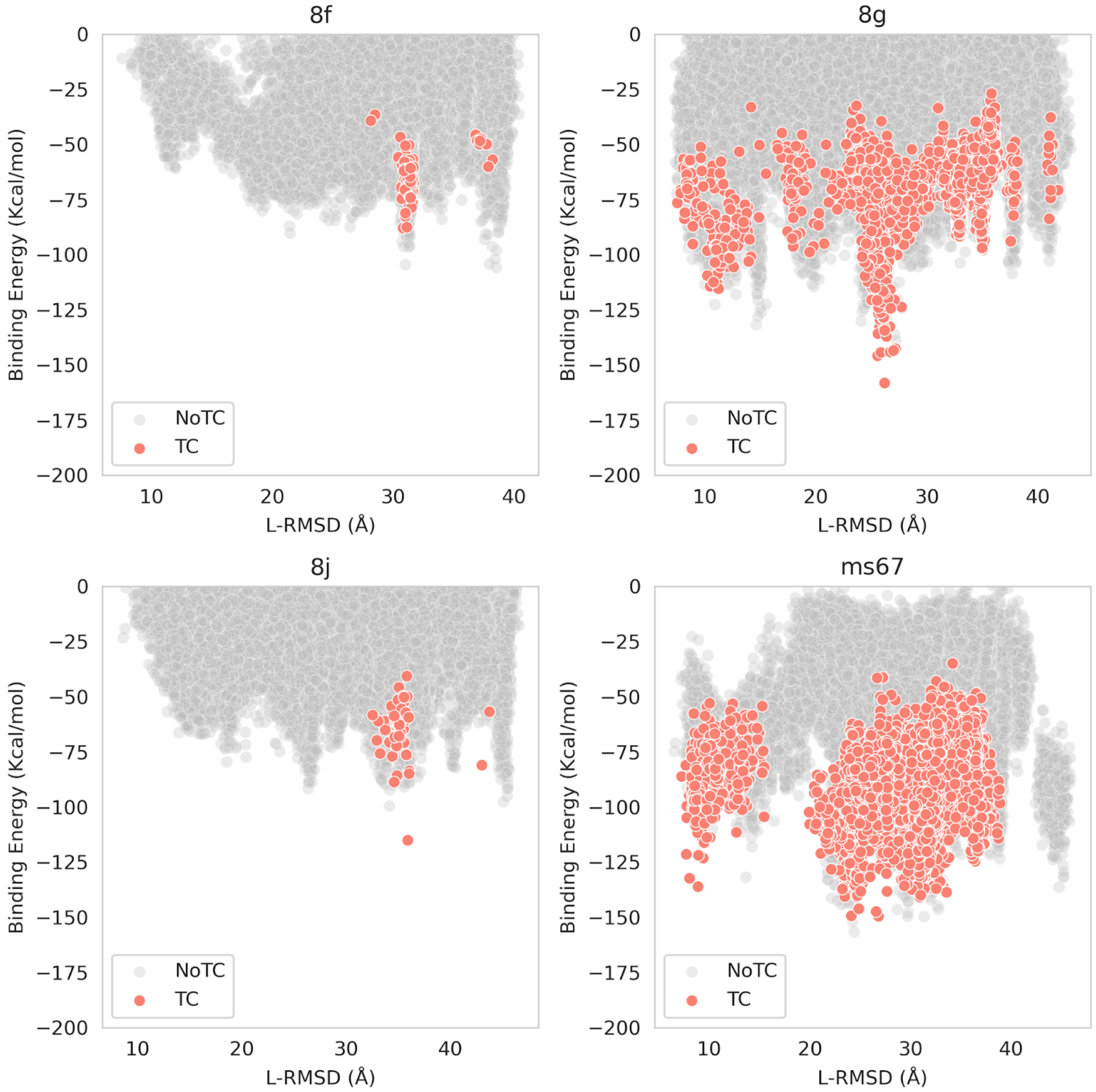


**Figure S18: Apo unbound binding energy TC landscapes**. Each point represents a system conformation accepted by PELE that forms a TC (orange) or not (gray).

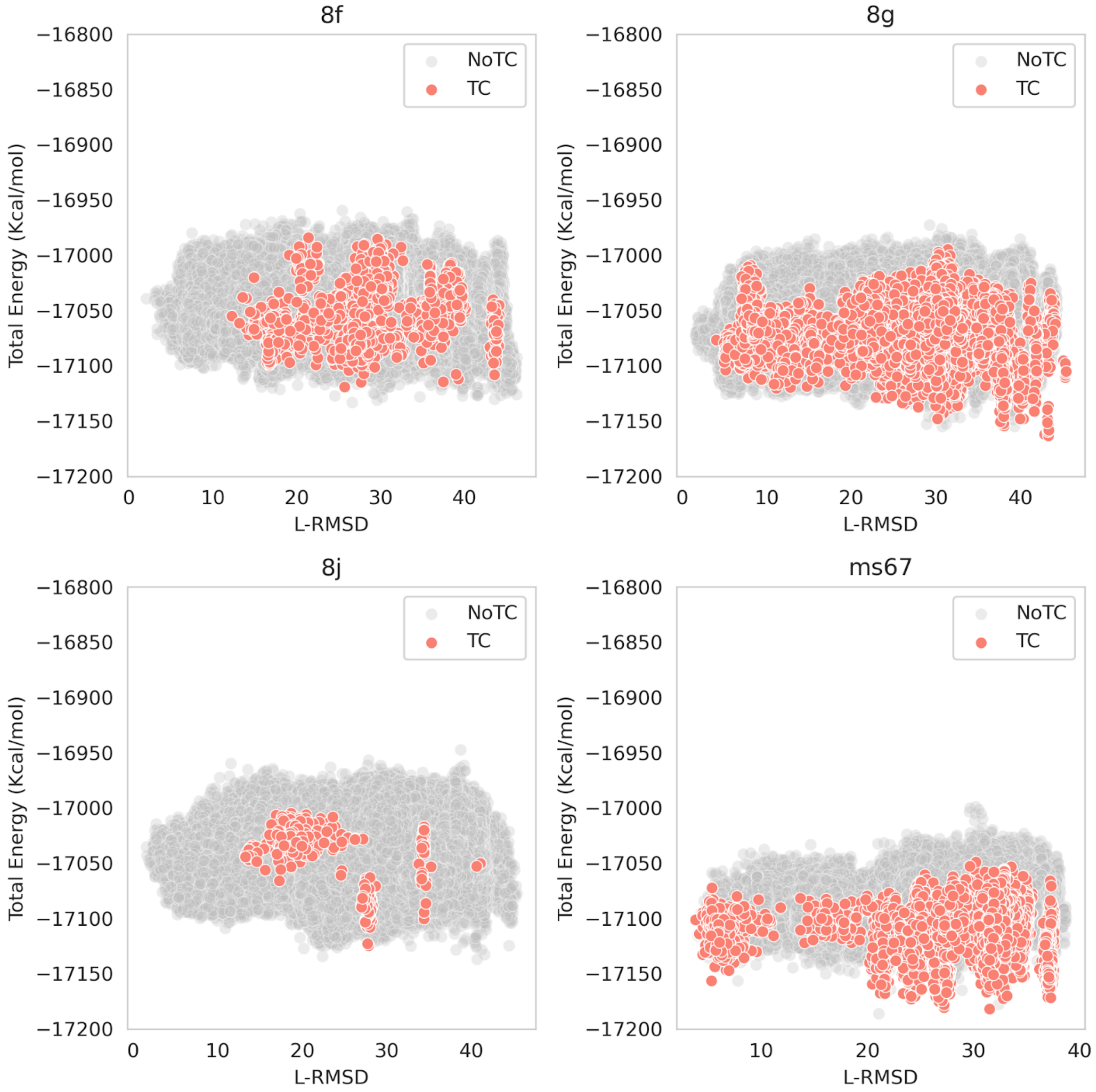


**Figure S19: Holo bound total energy TC landscapes**. Each point represents a system conformation accepted by PELE that forms a TC (orange) or not (gray).

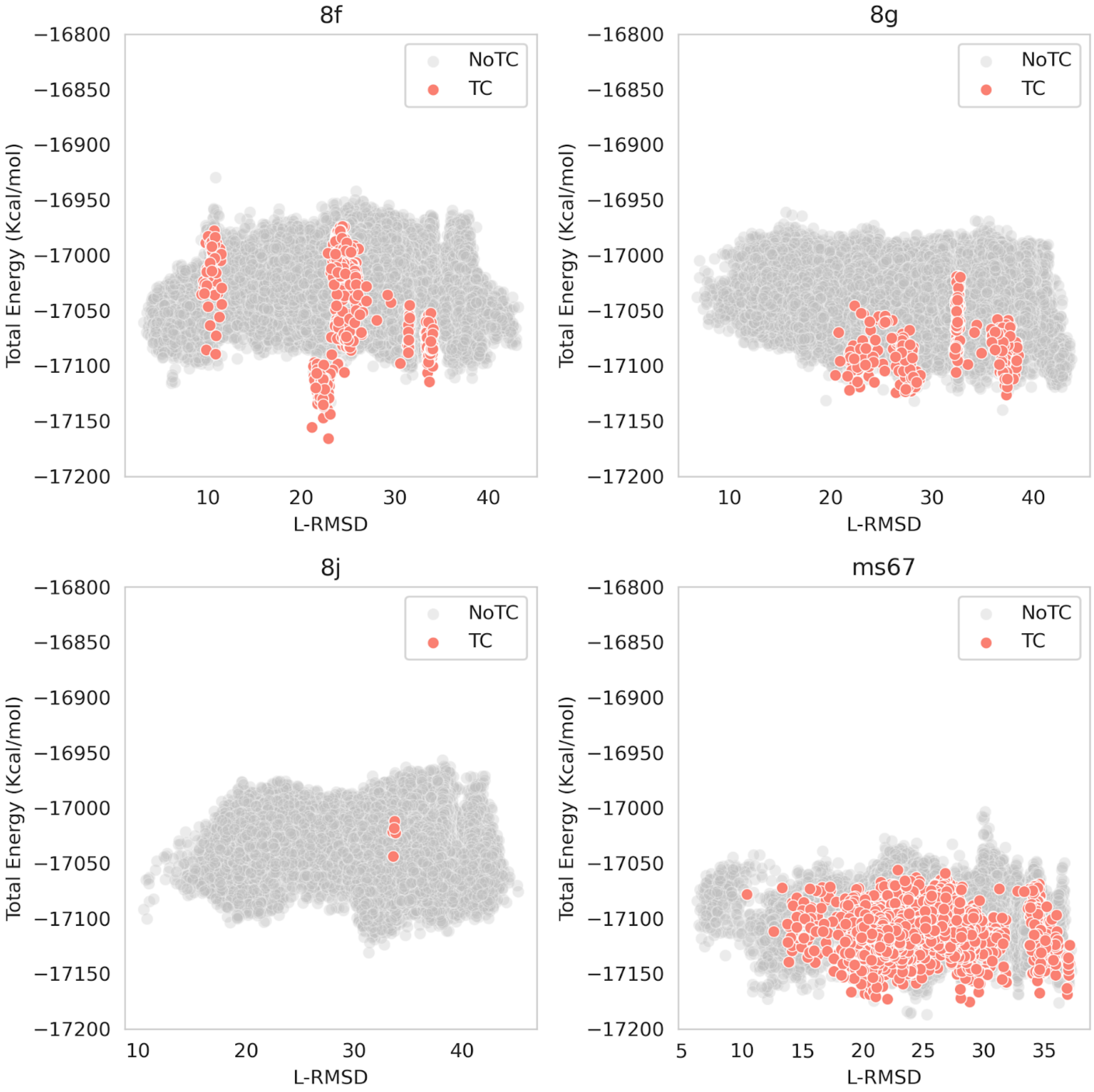


**Figure S20: Apo bound total energy TC landscapes**. Each point represents a system conformation accepted by PELE that forms a TC (orange) or not (gray).

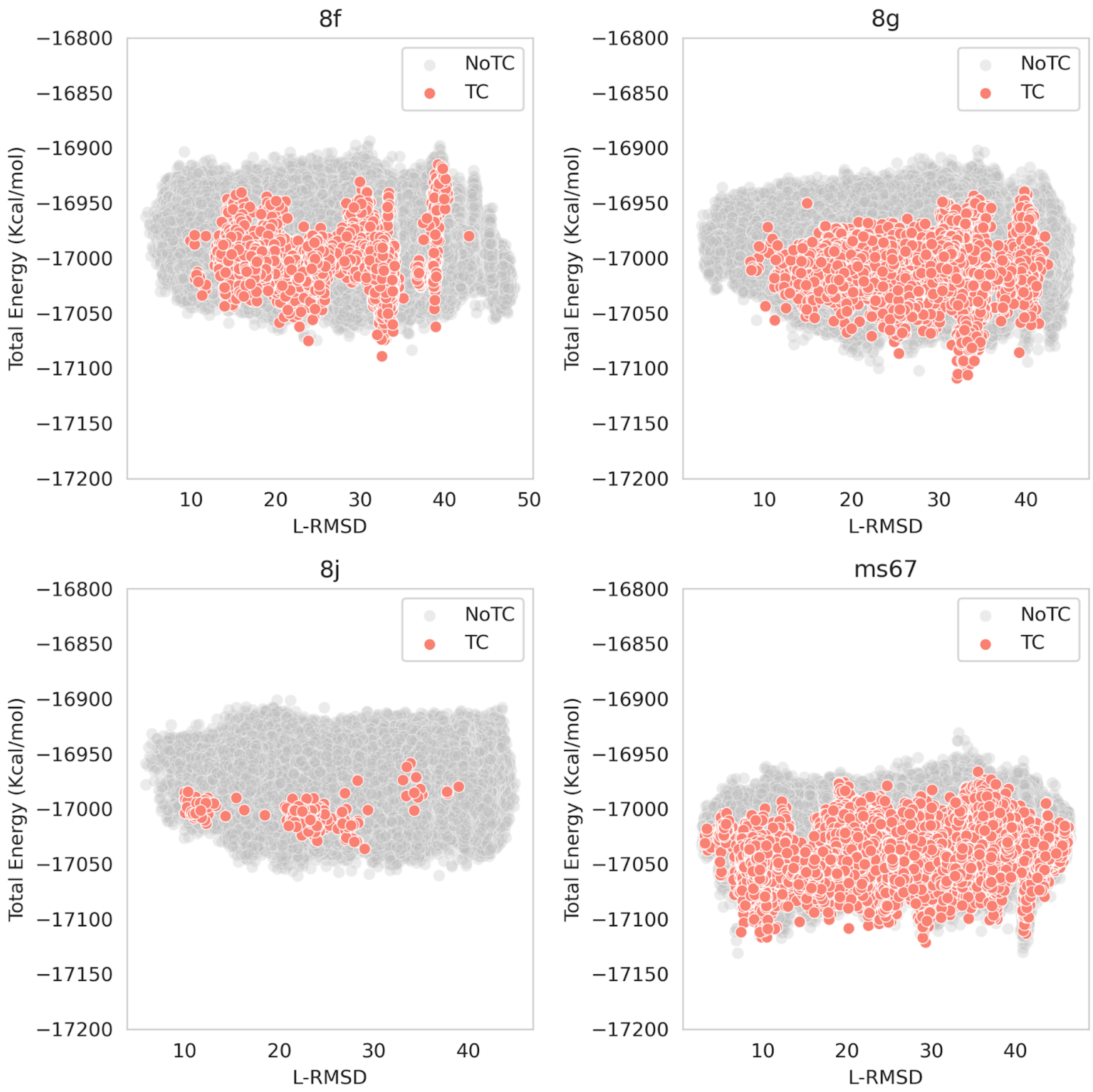


**Figure S21: Holo unbound total energy TC landscapes**. Each point represents a system conformation accepted by PELE that forms a TC (orange) or not (gray).

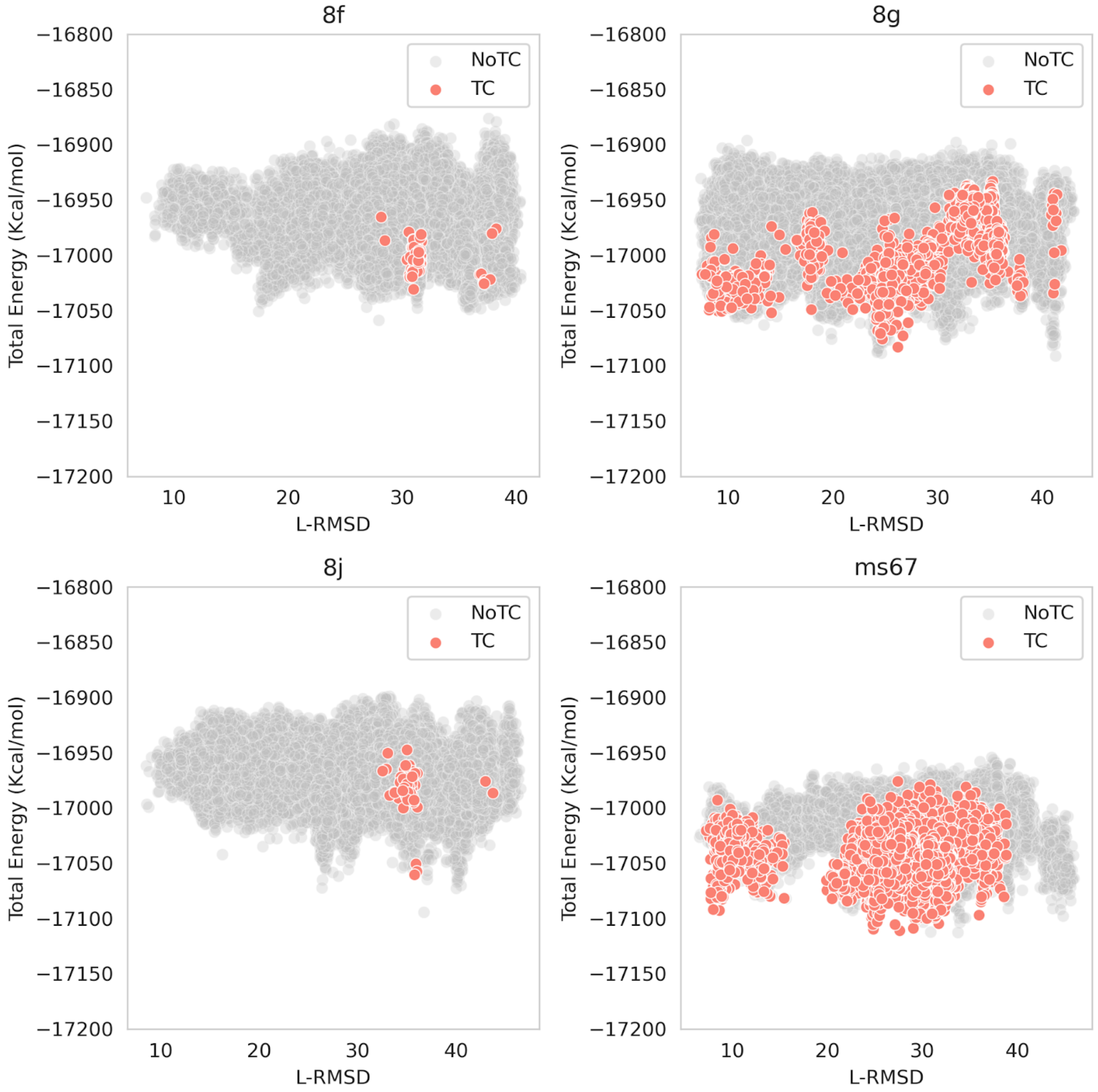


**Figure S22: Apo unbound total energy TC landscapes**. Each point represents a system conformation accepted by PELE that forms a TC (orange) or not (gray).

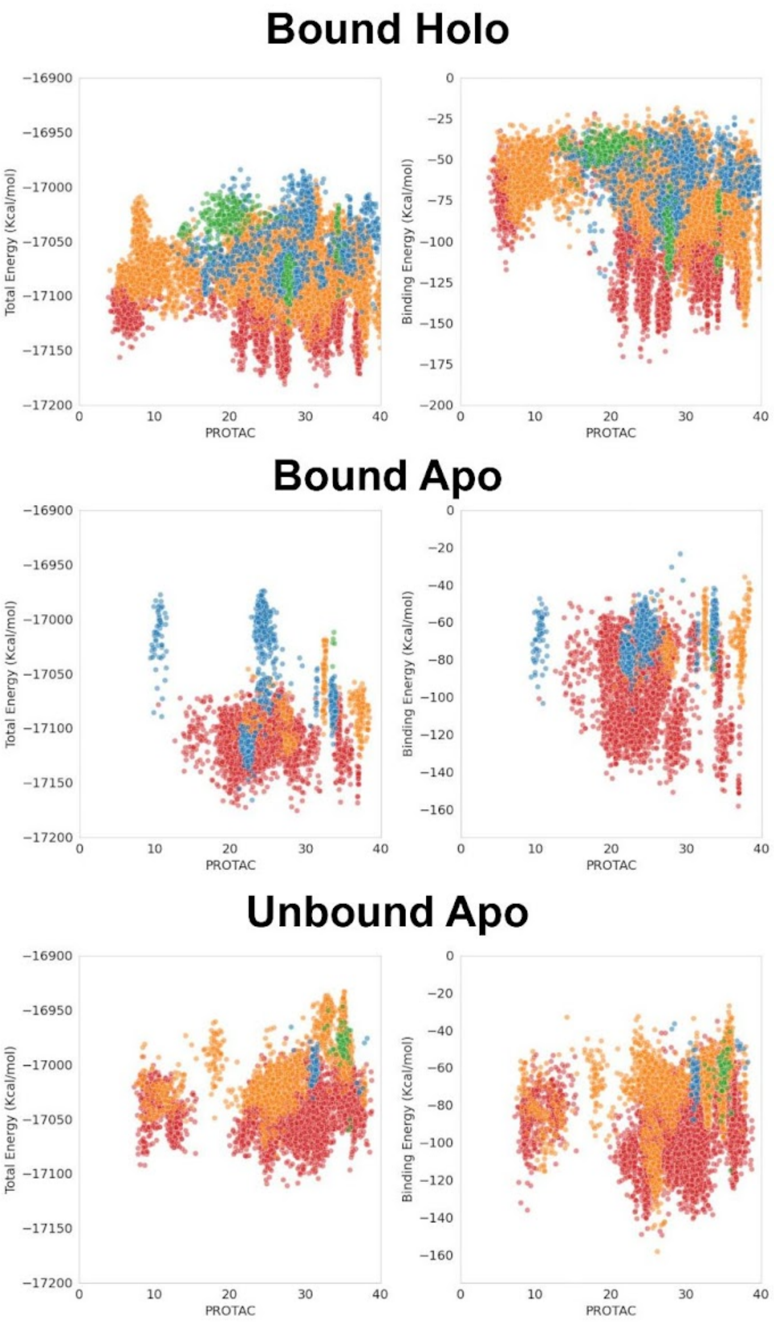


**Figure S23: Total and binding energies TC landscapes.** Each point represents a TC conformation found by PELE production for ms67 (red), 8g (orange), 8j (green), and 8f (blue). Refer to Figure 4A for the unbound holo landscapes.

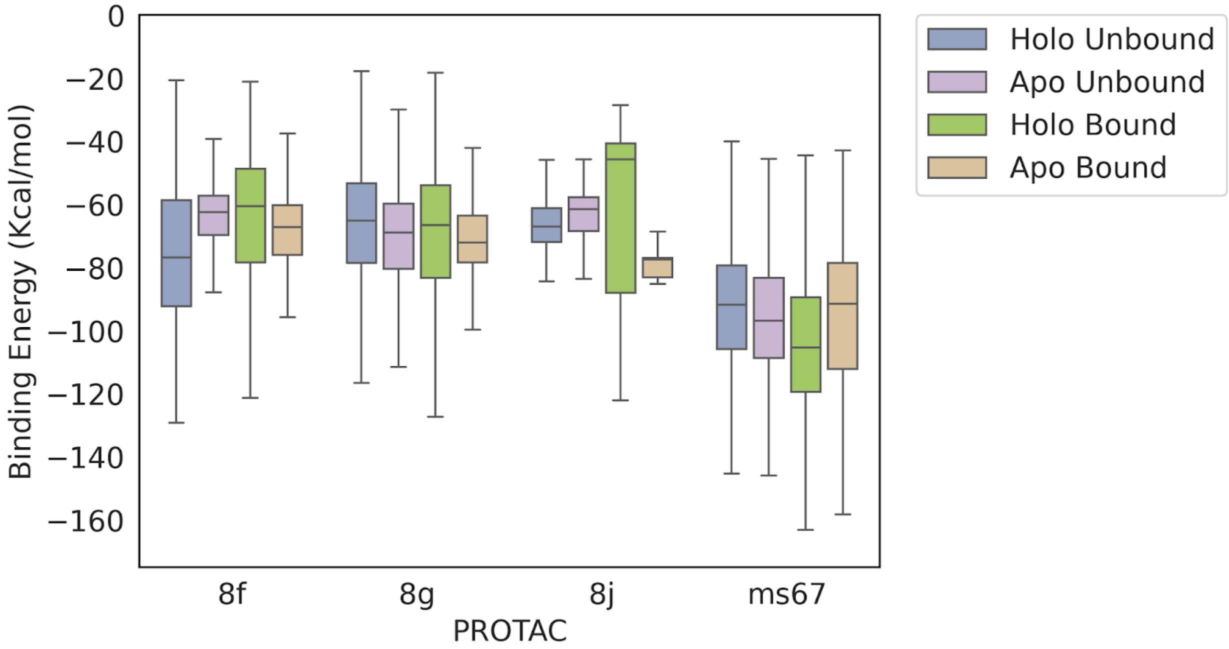


**Figure S24: Comparison of TC binding energy distributions for all studied modalities**.

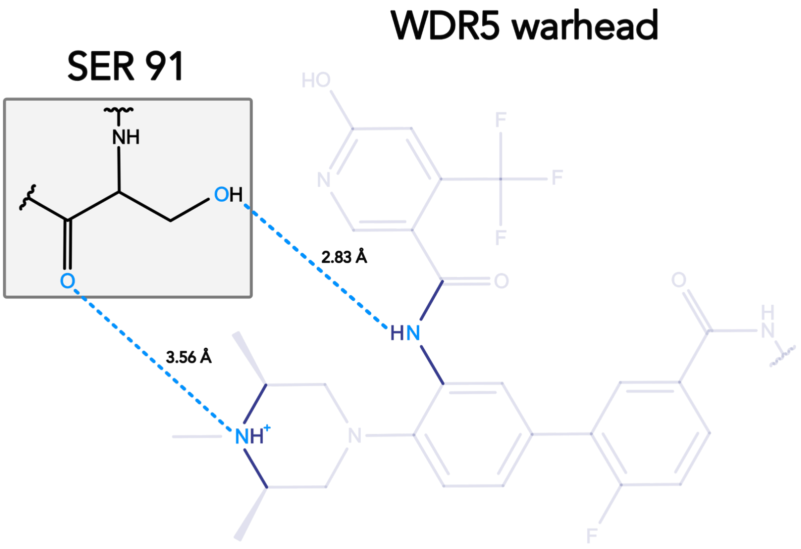


**Figure S25: Distance between the free warhead and its BS tracked during PELE simulations.** The dashed lines indicate the used distances.

### Supplementary Tables

**Table S1: Three point free warhead RMSD distributions of PROTAC sampling conformations relative to the crystal structure.** The PROTAC is docked at the VHL binding site, leaving the WDR5 warhead free. Each PROTAC shows as many RMSD distributions as its number of initial Glide docked poses. Distributions on the left correspond to the bound VHL conformation, while those on the right refer to the unbound conformation.

| PROTAC | # Initial clusters | | # PROTAC conformations with free warhead RMSD < 2 Å (#Total conformations sampled) | |
|---|---|---|---|---|
| | Bound | Unbound | Bound | Unbound |
| **ms67** | 2 | 4 | 8 (3,958) | 7 (7,263) |
| **8j** | 5 | 6 | 6 (9,358) | 6 (11,368) |
| **8g** | 5 | 6 | 10 (8,276) | 3 (9,108) |
| **8f** | 4 | 6 | 5 (5,371) | 5 (8,030) |

**Table S2: Cα-RMSD between bound and unbound crystals for VHL-WDR5 system.**

| Protein | Bound crystal (PROTAC) | Unbound crystal | Cα-RMSD (Å) |
|---|---|---|---|
| **VHL** | 7JTP (ms67) | 5NVX | 0.688 |
| | 8BB5 (8j) | | 0.690 |
| | 7Q2J (8g) | | 0.618 |
| | 8BB4 (8f) | | 0.560 |
| **WDR5** | 7JTP (ms67) | 4QL1 | 0.320 |
| | 8BB5 (8j) | | 0.333 |
| | 7Q2J (8g) | | 0.286 |
| | 8BB4 (8f) | | 0.595 |

**Table S3: Number of unique NN and PPD poses at different filtering stages for bound conformations.** Refer to **Table 1** for results using unbound conformations. Abbreviations: Equi - PELE Equilibration, Prod. - PELE Production.

| **PROTAC** | **8f** | | **8g** | | **8j** | | **ms67** | |
|---|---|---|---|---|---|---|---|---|
| **(NN / Unique PPD poses)** | **Holo** | **Apo** | **Holo** | **Apo** | **Holo** | **Apo** | **Holo** | **Apo** |
| **Step3 PPD** | 23 / 369,600 | 4 / 92,400 | 23 / 462,000 | 6 / 92,400 | 8 / 462,000 | 6 / 92,400 | 10 / 184,800 | 11 / 92,400 |
| **Step3 Filtered** | 23 / 34,362 | 11 / 8,588 | 23 / 43,892 | 6 / 8,588 | 8 / 43,648 | 6 / 8,588 | 10 / 22,488 | 4 / 8,588 |
| **Step 4 Fishing** | 20 / 1,855 | 3 / 811 | 17 / 3,359 | 3 / 1,025 | 4 / 4,485 | 1 / 1,298 | 7 / 578 | 7 / 417 |
| **Step 5 Equi.** | 85 / 1,855 | 24 / 811 | 133 / 3,359 | 15 / 1,025 | 85 / 4,485 | 17 / 1,298 | 15 / 578 | 8 / 417 |
| **Step 6 Prod.** | 23 / 164 | 7 / 43 | 38 / 442 | 3 / 72 | 16 / 159 | 0 / 42 | 10 / 148 | 2 / 63 |

**Table S4: Number of TC candidates and number of unique PPD poses that form a TC candidate at different filtering stages for bound conformations.** PROTACs are ordered from high to low ternary $K_D$ (see **Figure 2**). Refer to **Table 2** for results using unbound crystals.

| **PROTAC** | **8f** | | **8g** | | **8j** | | **ms67** | |
|---|---|---|---|---|---|---|---|---|
| **(Total TCs / Unique PPD poses)** | **Holo** | **Apo** | **Holo** | **Apo** | **Holo** | **Apo** | **Holo** | **Apo** |
| **Step 4 Fished** | 20,701 / 1,855 | 6,357 / 811 | 43,540 / 3,359 | 9,715 / 1,025 | 51,938 / 4,485 | 9,290 / 1,298 | 6,184 / 578 | 4637 / 417 |
| **Step 5 Equi.** | 25 / 12 | 14 / 6 | 397 / 87 | 21 / 10 | 15 / 13 | 1 / 1 | 862 / 68 | 648 / 47 |
| **Step 6 Prod.** | 1,318 / 30 | 682 / 7 | 10,471 / 89 | 365 / 9 | 410 / 10 | 5 / 1 | 5,551 / 57 | 3,159 / 33 |
